# Sample-level modeling of single-cell data at scale with *tinydenseR*

**DOI:** 10.1101/2025.11.26.690752

**Authors:** Pedro Milanez-Almeida, Daniela Schildknecht, Markus Linder, Saskia M. Brachmann, Andreas Weiss, Flavia Adler, Sofia Cardoni Lenticchia, Morgane Meistertzheim, Sophia Wild, Rachel Cuttat, Pushpa Jayaraman, Lang Ho Lee, Tanya Mulvey, Nadia Hassounah, Gina Crafts, David S. Quinn, Elena J. Orlando

**Affiliations:** Novartis Biomedical Research, Cambridge, MA 02139, USA; Novartis Biomedical Research, 4056 Basel, Switzerland

**Keywords:** single-cell data analysis, single-cell RNA sequencing (scRNA-seq), flow/mass/spectral cytometry, sample-level modeling, differential abundance analysis, differential expression analysis, single-cell atlas

## Abstract

Single-cell studies now routinely encompass hundreds of samples and millions of cells, offering unprecedented opportunities to link sample-level phenotypes with cellular and molecular states. However, current workflows often depend on cell-level inference and rigid clustering, which can distort significance and obscure subtle, continuous variation, in particular for complex experimental designs. Here, we present *tinydenseR*, a clustering-independent framework that enables robust, scalable, and statistically sensitive detection of differential cell state density, outperforming existing workflows in speed and memory usage. Technology-agnostic at its core, *tinydenseR* works seamlessly on scRNA-seq, flow, mass and spectral cytometry. Across synthetic benchmarks, a preclinical xenograft model, a publicly available COVID-19 study, two immuno-oncology trials and a multi-study atlas, *tinydenseR* uncovers disease and treatment history-associated effects, including subtle cell state heterogeneity, while embedding samples in a quantitative manner. Designed to accelerate discovery in clinical, preclinical, and translational research, the open-source package is available at GitHub.com/Novartis/tinydenseR.

## Introduction

The growing volume of single-cell data offers unprecedented opportunities to link sample-level phenotypes with cellular and molecular states. Studies now routinely profile hundreds of samples and millions of cells, enabling deeper insights into disease mechanisms and treatment responses [1-5]. However, this scale introduces significant computational challenges. Traditional workflows and data structures often fail to scale efficiently, creating bottlenecks in memory usage, processing time, and reproducibility when datasets reach atlas size [6-8].

Most current analysis strategies rely on clustering and cell-level inference as foundational steps for differential abundance (DA) and expression (DE) testing [9-11]. While clustering provides a convenient way to structure cellular diversity, it imposes rigid boundaries on inherently continuous biological processes. This simplification, combined with cell-level inference, can obscure subtle state changes and distort statistical significance, particularly within heterogeneous populations or complex experimental designs [12, 13]. Despite these drawbacks, clustering remains deeply embedded in single-cell workflows [14-17].

To improve efficiency and depth of analysis, many workflows employ landmark-based strategies, selecting representative cells to summarize large datasets [17-22]. These structure-aware approaches can preserve data geometry, rare populations, and local neighborhood structure more effectively than naive random subsampling. While landmark approaches reduce computational burden, they often remain tied to clustering or technology-specific assumptions, which can restrict flexibility and interpretability. More continuous methods, such as neighborhood count-based or embedding-driven analyses, aim to preserve biological resolution without rigid groupings. However, these approaches frequently depend on low-dimensional embeddings, which introduce uncertainty, or cannot generalize across technologies [19, 23-25].

In contrast, *tinydenseR* introduces a fuzzy set-based, technology-agnostic framework for sample-level modeling and quantitative embedding that avoids rigid clustering while preserving biological resolution (schematics in Fig. 1). By leveraging efficient data structures, *tinydenseR* scales to atlas-sized datasets and enables robust detection of differential cell state density across diverse technologies. We demonstrate its performance on synthetic benchmarks, a preclinical xenograft model, a publicly available COVID-19 study, two immuno-oncology trials, and a multi-study atlas, uncovering treatment-associated effects, including subtle cell state heterogeneity, while outperforming popular current workflows.

**Figure 1.**
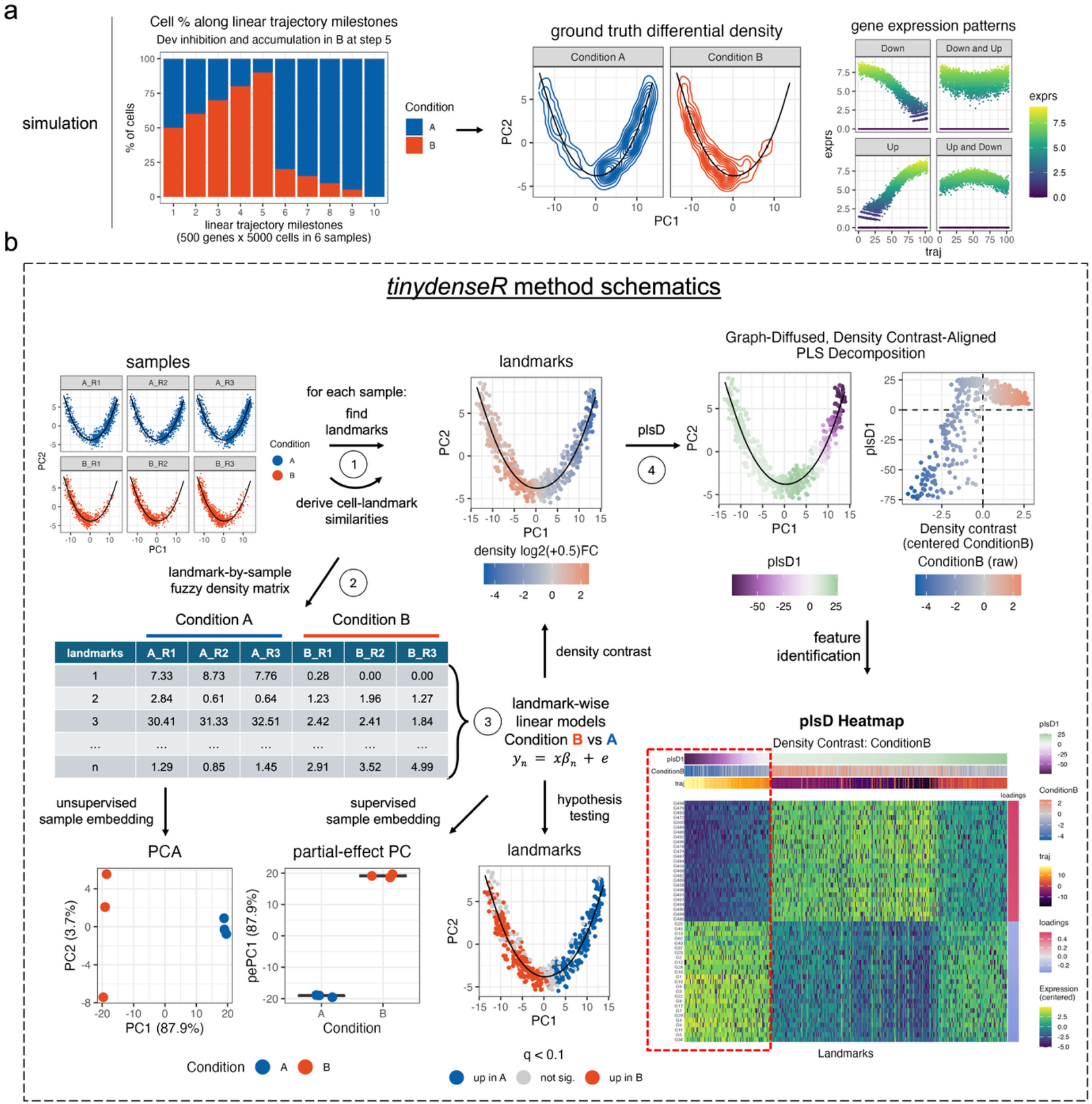
Synthetic trajectory benchmark and overview of the *tinydenseR* workflow. (a) Synthetic scRNA-seq trajectory dataset used to illustrate the method. Left, proportion of cells along discrete milestones of a linear trajectory across two conditions (A and B), showing a condition-dependent redistribution of cells along the trajectory. Middle, ground-truth differential density displayed on the trajectory manifold for each condition. Right, example gene-expression patterns defined on the same trajectory (traj, x-axis), illustrating that molecular programs can vary along the trajectory and need not coincide with discrete boundaries. The synthetic benchmark contains two conditions with three samples per condition. (b) Schematic of the *tinydenseR* workflow. Cells from each sample are summarized by a shared set of landmark cells. For each sample, cell-to-landmark connection strengths are computed and aggregated to form a landmark-by-sample fuzzy density matrix. This matrix is used for sample-level modeling of landmark densities, yielding landmark-wise estimates of differential density (that is, the density-contrast log2 fold changes) and associated hypothesis testing. The same density matrix also supports sample embedding, including unsupervised PCA and supervised partial-effect principal component (pePC) projections, the latter isolating the modeled effect of interest. To aid interpretation of the density contrast, *tinydenseR* further applies graph-diffused, density-contrast-aligned partial least squares decomposition (plsD) to identify expression programs aligned with the inferred density shift, visualized as landmark scores, top features, and a feature-by-landmark heatmap, where rows show centered expression of top 25 features associated with the component in each direction. For formal DE analysis of selected populations, sample-level pseudobulks are constructed using landmark connection-strength weights, as detailed in Materials and Methods (not shown).

## Results

### The tinydenseR Algorithm

To avoid hard cluster boundaries, *tinydenseR* represents each sample by its UMAP-derived fuzzy density around a shared set of landmark cells. This yields a novel sample representation in *tinydenseR* – the landmark-by-sample fuzzy density matrix *Y* – in which individual cells contribute fractionally to multiple nearby landmarks according to their fuzzy graph affinities, thereby preserving continuous cell-state variation rather than forcing cells into discrete clusters. *tinydenseR* then performs sample-level modeling on the size-factor-normalized and log2-transformed matrix *Y*, using standard design matrices with optional covariates and blocking, to formally test for associations between cell-state density and sample-level variables without requiring that the relevant states coincide with a particular clustering solution (schematic in Fig. 1; details in Materials and Methods). We refer to this as differential cell-state density analysis, and to the corresponding modeled log2-fold change around each landmark as the density contrast.

To summarize the sample-level signal associated with a given density contrast while retaining the covariate structure of the model, *tinydenseR* provides a novel contrast-specific sample embedding we term pePC (partial-effect Principal Component projections). pePC is derived from the model’s partial fitted values for the density contrast of interest, so that the resulting embedding isolates the modeled change attributable to that effect rather than raw confounded variation. As a result, separation in pePC space reflects alignment with the density contrast and can be used to visualize continuous gradients or discrete separations across samples under complex designs, including settings with covariates or repeated measures. pePC also provides a quantitative summary of how much of the total variation in the landmark-by-sample matrix is captured by the modeled effect (schematic in Fig. 1; details in Materials and Methods).

Naturally, *tinydenseR* also returns unsupervised sample embeddings of the landmark-by-sample density matrix (that is, PCA and diffusion-map trajectory), which summarize dominant axes of sample-to-sample variation without specifying a covariate of interest. The unsupervised sample embeddings together with the pePC supervised embedding for a density contrast jointly help surface unexplained structure: if unsupervised embeddings reveal separation that is not aligned with the pePC axis (or if pePC explains only a small fraction of total variance), this suggests that additional biological or technical factors – measured or unmeasured – remain to be modeled.

To help interpret the features driving the density contrast of interest, *tinydenseR* provides a multivariate decomposition that we term plsD (graph-diffused, density contrast-aligned <u>p</u>artial least <u>s</u>quare <u>D</u>ecomposition). Rather than testing each gene or marker independently, plsD identifies latent axes of graph-smoothed expression that are maximally aligned with the landmark-level direction of change in density. That means, plsD highlights genes or markers associated with differential cell state density due to differential abundance, expression, or both, within the same landmark framework used for density modeling. Thus, plsD provides interpretable exploratory landmark-maps of expression programs that are most strongly associated with the observed cell state density shift (schematic in Fig. 1; details in Materials and Methods).

In addition, *tinydenseR* supports formal sample-level inference for selected populations through pseudobulk differential expression analysis. Two modes are available: design mode, which tests for differential expression across groups of samples within a selected subset of landmarks (for example, classical monocytes before versus after infection), and marker mode, which compares subsets of landmarks within each sample to identify features associated with particular cell states (for example, clusters of resting vs activated T cells after treatment). In either mode, inference is based on standard linear modeling, but *tinydenseR* differs from conventional pseudobulk workflows in how the pseudobulks are constructed: expression is aggregated fractionally as a connection strength-weighted sum over landmark neighborhoods rather than by hard cluster or cell-type membership. This affinity-weighted aggregation integrates expression over the local neighborhood structure captured by the cell– landmark graph, yielding a manifold-informed pseudobulk representation while retaining samples as the unit of inference. These two pseudobulk DE modes complement the subsetting-free, exploratory plsD framework by providing formal gene-level statistical testing with samples as biological replicates. In the next sections, we evaluate *tinydenseR* across a range of datasets, including synthetic data with known ground truth as well as real-world clinical, preclinical, and translational studies, and compare its performance with other widely used R packages for related tasks.

### Detection of Differential Cell State Density in Synthetic Benchmarks

To start evaluating the statistical behavior and robustness of *tinydenseR*, we applied it to three synthetic datasets designed to simulate key single-cell analysis tasks across different technologies. The first dataset simulated differential continuous trajectory patterns in single-cell RNA-sequencing (scRNA-seq) and was generated along a single linear trajectory with 10 discrete milestones, comprising 5,000 cells and 500 genes (Fig. 1a). Condition effects were introduced by systematically varying the proportion of cells assigned to conditions A and B at each milestone, thereby creating differential cell-state abundance along the trajectory while preserving the underlying continuous structure. Cells were further labeled with replicate identifiers, yielding sample labels defined by condition and replicate, as described in Materials and Methods. *tinydenseR* recovered trajectory-associated cell-state changes, embedded samples along the variable of interest and uncovered gene expression programs associated with changes in cell density as expected, illustrating its ability to model continuous cellular and molecular variation without relying on hard cluster boundaries (Fig. 1).

The second synthetic dataset mimicked a flow cytometry differential abundance experiment. Three settings were simulated, corresponding to two-fold depletion of a large (50% to 25%), small (5% to 2.5%), or very small (0.5% to 0.25%) target cell population across treatment groups, with six samples per group, approximately 50,000 cells per sample, and five markers in each setting. Marker-specific batch effects and cell-intrinsic expression heterogeneity were introduced independently of abundance changes, such that differential abundance of the target population was the sole systematic treatment effect. *tinydenseR* accurately detected all three abundance shifts, including the very small population change (Fig. S1). Of note, negative loadings in plsD1 for markers of the target population reflected the ground truth target depletion across all three settings, while marker-mode pseudobulk DE – that is, comparing the cluster of cells most affected in the density contrast against all other clusters – correctly identified markers of the target population (Fig. S1a-c, features).

The third synthetic dataset mimicked a flow cytometry differential expression experiment. In this simulation, the target population was fixed at 5% of total cells in all samples, and differential signal was introduced exclusively through activation-dependent upregulation of Marker2 within target cells. Three effect sizes were simulated, corresponding to shifts in the log-mean of Marker2 expression from 0 to 2, 0 to 1, or 0 to 0.5, with six samples per group, approximately 50,000 cells per sample, and five markers in each setting. Because target-cell abundance was held constant across groups, systematic treatment differences arose from within-population expression changes rather than compositional shifts. *tinydenseR* successfully identified all three expression shifts, including the smallest simulated change (Fig. S2). Positive loadings in plsD1 for Marker2 reflected the ground truth target activation across all three settings, while design-mode pseudobulk DE – that is, comparing treatment vs baseline within the cluster of cells most affected by the density contrast – correctly identified Marker2 as most differentially expressed (Fig. S2a-c, features).

Analysis of the cytometry datasets with *diffcyt*, which implements statistical methods for differential discovery analyses in high-dimensional cytometry data [15], also detected the expected differences associated with depletion and activation in both simulations (Fig. S3). However, *tinydenseR* was able to correctly identify affected markers in an unbiased manner, whereas *diffcyt* requires, by design, manual distinction of lineage and functional markers for clustering and DE analysis, respectively. This reliance on prior knowledge can restrict the discovery of unexpected treatment effects.

Across the three synthetic benchmark settings, permutation analyses were dominated by relabelings with either no q-significant discoveries or only weak minimum q-values, whereas the strongest signals occurred for the observed labels and their exact complements (Fig. S4). These results suggest limited spurious discovery under most arbitrary relabelings in the benchmark settings considered, while also showing high sensitivity when the labels are strongly aligned with the underlying sample structure. Consistent with this interpretation, *tinydenseR* maintained high sensitivity and low estimated false discovery rate across the benchmark settings, even in scenarios where the modeled effect explained only a small fraction of the total variance in the density matrix (Fig S1c and S2c). Together, these results validate *tinydenseR*’s capacity to handle diverse single-cell analysis tasks while also enabling unbiased, unexpected and clustering-free discoveries.

To assess whether landmark-level integration preserves biological structure relative to cell-level integration, we compared *tinydenseR* with Harmony on all cells using the Luecken et al. Immune_ALL_human benchmark dataset (33,506 cells; 10 batches). Landmark-level integration with *tinydenseR* matched or slightly exceeded cell-level Harmony on four of six integration-quality metrics, including all three biological-conservation metrics (MCC for cell-type label transfer, cell-type silhouette width and graph connectivity), whereas cell-level Harmony performed better on the two neighborhood-based batch-mixing metrics (kBET and batch entropy) (Fig. S5a-c). Intriguingly, the UMAP layouts differed qualitatively in progenitor/HSPC-related regions, with landmark-level integration appearing more elongated and continuous, while cell-level integration somewhat compacted cells together in the same regions (Fig. S5a-b). These results indicate that compression to landmarks did not measurably impair preservation of major biological structure in this dataset.

### Application to Preclinical and Clinical Real-World Datasets

Next, we investigated treatment-induced transcriptional changes in a preclinical setting with *tinydenseR*. For this, we analyzed a xenograft model dataset generated by scRNA-seq. KRAS(G12D) mutant SW1990 pancreatic cancer cells, either wild-type (WT) for *HRAS* and *NRAS* (H/NRAS) or engineered with a H/NRAS double knockout (KO), were implanted into mice that were subsequently treated for seven days with either a vehicle or the KRAS(G12D) inhibitor MRTX1133 (Fig. S6a; see Materials and Methods for experimental details). This design enabled comparison of the treatment effect within each genetic background.

*tinydenseR* revealed distinct shifts in the density of transcriptionally defined cell states, with notable differences between WT and H/NRAS KO tumors. For example, KO cells from mice treated with KRAS(G12D) inhibitor or vehicle separated into two distinct regions of the manifold with clear differences along the treatment-versus-vehicle density contrast, indicating a strong treatment-associated effect in KO cells (Fig. S6b, KO). In contrast, WT cells from mice treated with KRAS(G12D) inhibitor or vehicle were distributed within a broader region of the manifold with local peaks and valleys along the WT density contrast, suggesting a more heterogeneous treatment-associated response in WT cells (Fig. S6b, WT). Supervised quantitative embedding of samples along each density contrast further showed that approximately three times as much variance in the density matrix was explained by treatment in KO compared with WT cells (28.3% versus 8.2%, respectively; Fig. S6c), corroborating that KO cells respond to treatment more homogeneously than WT cells at the level of transcriptionally defined cell states.

Exploratory gene set enrichment analysis of plsD loadings for the KO density contrast was consistent with treatment-associated changes in KRAS-related and cell-cycle-associated transcriptional programs (Fig. S6d). In WT cells, analogous pathway structure was less prominent on plsD1, whereas plsD2 highlighted an additional treatment-associated axis involving a subset of WT landmarks with a potential shift in PI3K/AKT/MTOR signaling that was not prominent in KO plsD1 (Fig. S6e–f). The trends observed with plsD2 were supported by inference with design mode pseudobulk DE using the WT treatment contrast on the subset of landmarks at least 1 SD from the mean WT plsD2 score and at least 0.5 log2-fold change in density contrast in the same direction. These results highlight the utility of *tinydenseR* in dissecting complex treatment responses in preclinical models by quantitatively modeling compositional and functional heterogeneity. In addition, exploratory analysis with plsD helped generate hypotheses about transcriptional programs that might help explain differences in response to KRAS(G12D) inhibition between H/NRAS KO and WT tumors.

Next, we analyzed the COMBAT COVID-19 scRNA-seq dataset [26], which contains information about clinically relevant and heterogeneous phenotypes. After restricting the analysis to healthy volunteers (HV) and COVID-19 samples classified as mild, severe or critical, and controlling for sex and age, *tinydenseR* was able to detect cell-state heterogeneity and quantitatively embed samples along disease status, severity and time since onset of worsening symptoms. Importantly, time since onset of worsening symptoms differed across severity groups (Fig. S7a), raising the possibility that apparent severity-associated differences in PBMC immune cell states could be influenced in part by sampling time. To help reduce this potential confounding, and to allow for non-linear effects of disease duration, time since onset was binarized into early and late strata among infected samples, yielding contrasts for: early infection versus healthy; late versus early infection; and early critical versus early mild disease (Fig. S7a–d). These analyses revealed broad landmark-level density shifts across major immune compartments, including monocyte/dendritic cell, T/NK and B/plasmablast-rich regions of the manifold (Fig. S7b–c), while supervised quantitative embedding separated samples along the modeled contrasts, with the largest effect size observed for early COVID versus healthy (10.4% of variance explained by pePC1), followed by late versus early infection (3.7%) and early critical versus early mild disease (1.7%) (Fig. S7d).

To further interpret the early critical-versus-mild contrast, we applied plsD to the corresponding density contrast. The leading plsD component highlighted a monocyte-centered transcriptional program consistent with a dysregulated inflammatory state in early critical disease, with positive loadings associated with neutrophil degranulation-related structure and negative loadings associated with interferon alpha and interferon gamma response programs (Fig. S8). The genes contributing most strongly to this component included inflammatory myeloid features such as *S100A12, CLU, VCAN* and *CTSD* on one side, and interferon/HLA-associated features such as *HLA-DPA1, HLA-DPB1, IFITM1, IFIT2 and IFIT1*, on the other (Fig. S8b-c). These findings are consistent with a shift from interferon-responsive classical monocytes in early mild disease toward a more inflammatory, dysregulated myeloid program in early critical disease, in line with previous reports of myeloid dysregulation in severe COVID-19 [27].

A second plsD component further resolved heterogeneity within the same contrast; notably, plsD2 scores were most extreme within two very small subsets of landmarks (Fig. S9a). Analysis of plsD2 loadings revealed the immature/progenitor-like cell-associated genes *CD34, SMIM24, CYTL1, CPA3, CPXM1* and *CRHBP* to be highly expressed in landmarks with the particularly high plsD2 scores, while the plasmacytoid dendritic cell (pDC)-associated genes *CLEC4C, LILRA4* and *TPM2* were highly expressed in landmarks with particularly low plsD2 scores (Fig. S9b-c). Furthermore, the natural killer (NK) cell-associated genes *NKG7, PRF1, CST7, GZMB, CCL5, GZMH* and *GZMA* were among the 100 genes with the most negative loadings (respective ranks: 43, 44, 57, 13, 41, 33 and 42). These findings are also consistent with reports that severe COVID-19 is accompanied by emergency myelopoiesis and impaired antiviral innate immunity, including expansion of proliferative progenitor/myeloid states and reduced or dysfunctional pDC and NK cell programs [5, 27], suggesting that plsD2 may capture a secondary severity-associated axis of hematopoietic stress beyond the dominant dysregulated monocyte program highlighted by plsD1. Together, these results illustrate how *tinydenseR* can combine sample-level modeling with exploratory decomposition to resolve multiple layers of biological heterogeneity within clinically relevant disease contrasts associated with rare cell states.

We then applied *tinydenseR* to a single-cell dataset from a phase 1 immuno-oncology clinical trial evaluating NIZ985, a recombinant heterodimer of IL-15 and IL-15 receptor α, in combination with PDR001, an anti-PD-1 monoclonal antibody, in adults with metastatic cancers. The dataset included longitudinal 12-marker flow cytometry profiles from peripheral blood mononuclear cells (PBMC) from 11 patients, with samples collected at baseline (Cycle 1 Day 1, C1D1) and at multiple post-treatment timepoints (C1D3, C1D4, C1D8, C1D15 and C1D17; Fig. S10a). We focused on analysis of the CD8 T cell compartment here.

*tinydenseR* identified treatment-associated shifts in the density of CD8 T cell states across timepoints relative to baseline (Fig. S10b). Supervised quantitative embedding of the density matrix along visit/timepoint showed that pePC1 separated C1D1 and C1D4 samples most clearly, with pePC1 explaining 9.1% of the total variance in the density matrix, indicating that the strongest modeled visit-associated effect occurred at C1D4 (Fig. S10c). By contrast, unsupervised PCA retained substantial structure associated with batch and subject ID rather than isolating the treatment-time effect (Fig. S10c). These results underscore *tinydenseR*’s ability to appropriately partial-out repeated-measures and batch effects in complex longitudinal designs. Exploratory plsD analysis of the C1D4 versus C1D1 density contrast indicated that the dominant treatment-associated program was enriched for markers of proliferation and activation, with the strongest positive loadings for Ki67, CD38 and HLA-DR (Fig. S10d). The C1D4-enriched end of the landmark-level heatmap also showed generally lower CCR7 and low-to-intermediate CD45RA expression, albeit with appreciable heterogeneity across landmarks. Together, these results highlight *tinydenseR*’s ability to resolve transient, treatment-associated activation states within CD8 T cells while modeling samples as the unit of inference in a longitudinal clinical setting.

To further evaluate *tinydenseR* in a clinical trial context, we analyzed scRNA-seq data from a phase 1 trial of PHE885, a rapidly manufactured BCMA-targeting CAR T cell therapy, in 25 patients with newly diagnosed multiple myeloma (NDMM) and 42 patients with relapsed or refractory multiple myeloma (RRMM). The dataset included 321 samples and 1.76 million cells post-quality control (median 4,725 cells/sample) from leukapheresis, final product, and longitudinal PBMC and bone marrow mononuclear cells (BMMC; day 28, month 3, month 6; Fig. S11). *tinydenseR* enabled multi-level modeling of abundance shifts across timepoints and compartments, revealing changes associated with treatment and disease history in cellular composition and transcriptional states.

At screening in PBMC, landmark annotation showed that the manifold was organized primarily into T/NK, monocyte/dendritic-cell, and B/plasmablast/plasma-cell rich regions (Fig. S12a), and the fitted NDMM versus RRMM density contrast localized cohort-associated differences across these compartments, with prominent positive and negative signals mixed in T/NK-rich regions and in monocyte-rich regions (Fig. S12b). Supervised quantitative embedding separated NDMM and RRMM samples along pePC1, which explained 7.1% of the total variance in the density matrix (Fig. S12c). The corresponding density contrast and plsD1 loadings suggested relatively greater representation of early activated transitional T cell structure in NDMM, with positive loadings for genes such as *SBDS, JUND, BTG1, CD69*, and *PIK3IP1*, whereas the opposite side of the contrast was characterized by effector/memory-like cytotoxic T cell-associated genes including *GZMA, IL32, PTPRCAP* and *PRF1* (Fig. S12b,d–e). These results are consistent with disease- and prior-treatment-associated systemic immune remodeling in RRMM relative to NDMM [28].

During manufacturing, *tinydenseR* identified a strong shift between leukapheresis (Aph) and final product (FP), with opposing directions of the density contrast localizing predominantly to T/NK-rich (higher density) versus monocyte/DC-rich (lower density) regions of the manifold (Fig. S13a-b). Supervised quantitative embedding separated FP and Aph samples along pePC1, which explained 71.3% of the total variance in the density matrix (Fig. S13c), while plsD1 further indicated that the FP-associated program was enriched for genes involved in proteostasis/chaperones (*HSP90AA1, HSP90AB1, HSPA8, HSPD1* and *HSPE1*), RNA/translation-related machinery (*NPM1, RAN, EIF5A, NCL* and *SERBP1*), metabolic/biosynthetic activity (*LDHA, ENO1, PGAM1* and *CHCHD2*), and cytoskeletal remodeling (*TUBA1B* and *TUBB*), whereas negative loadings included the T cell quiescence-associated transcription factor *KLF2* and monocyte-associated genes such as *LYZ, VCAN* and *CD14* (Fig. S13d–e). Together, these results are consistent with enrichment of biosynthetically active, non-quiescent T cell states and depletion of non-T cell contaminants during rapid CAR T manufacturing [29].

In the bone marrow, landmark-level density contrasts revealed heterogeneous changes across the manifold at day 28, with weaker residual structure at month 3 and little remaining signal by month 6 (Fig. 2a-b). Supervised quantitative embedding blocking for intra-subject correlation and controlling for disease history showed that day 28 was the timepoint most clearly separated from screening along pePC1, which explained 6% of the total variance in the BMMC density matrix (Fig. 2c). Exploratory plsD analysis of the day 28 versus screening contrast resolved at least two transcriptional programs: plsD1 was associated with inflammatory-monocyte (positive loadings) and plasma-cell-related structure (negative loadings), whereas plsD2 highlighted an additional memory-T-cell-associated axis (positive loadings) opposed to additional plasma-cell structure (negative loadings; Fig. 2d–e). Finally, CAR construct expression across BMMC landmarks was most strongly associated with increased abundance at day 28, was attenuated at month 3, and was largely absent by month 6 (Fig. 2f), consistent with peak CAR T-cell expansion in bone marrow at day 28 followed by contraction over time. Together, these patterns are consistent with a day 28 bone marrow response marked by reduced plasma-cell-associated structure and increased inflammatory monocyte and CAR-associated memory T cell structure, in line with the expected pharmacodynamic effects of a BCMA-directed CAR T cell therapy.

**Figure 2.**
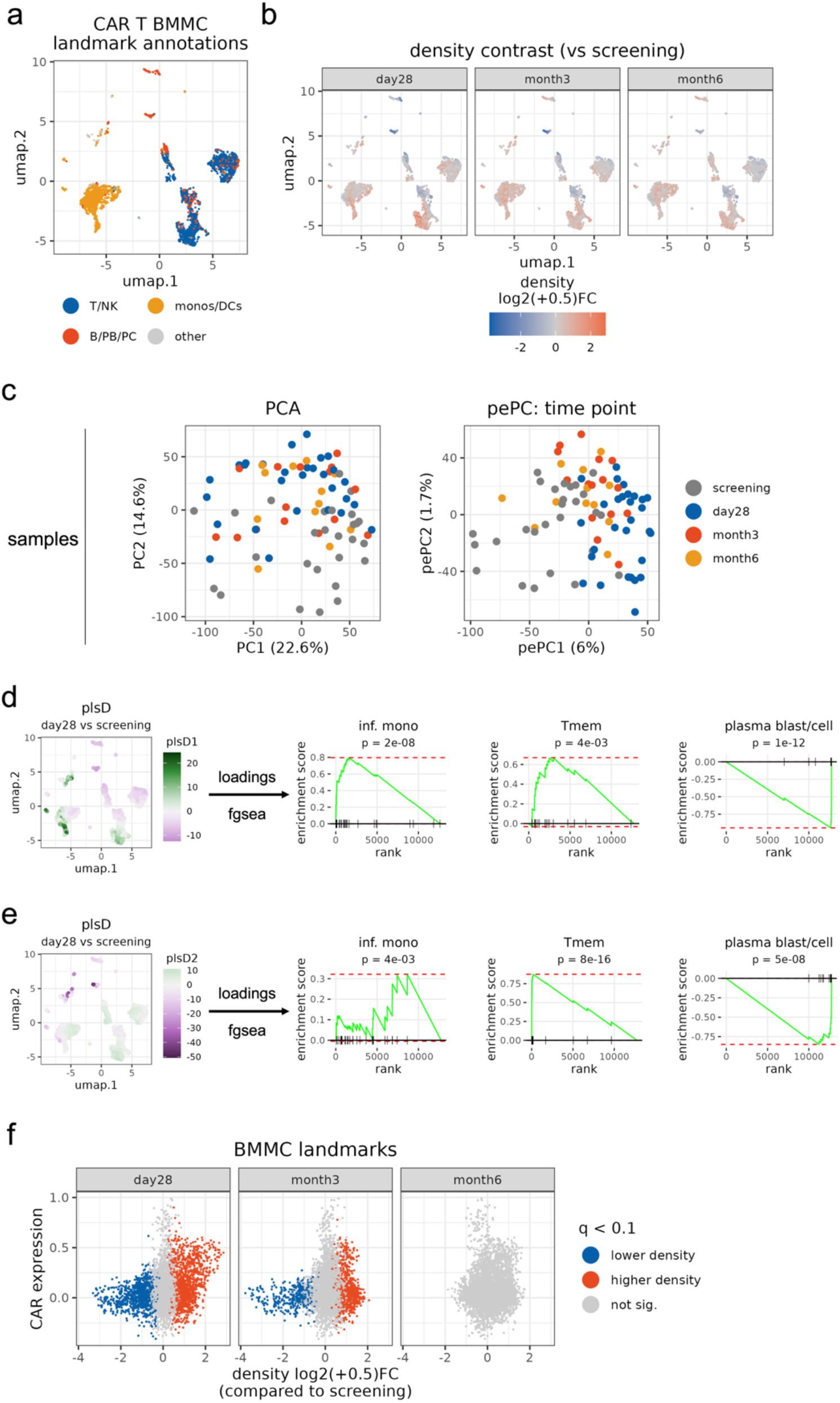
Modeling BMMC single-cell data from multiple myeloma patients after BCMA-targeting CAR T-cell infusion using *tinydenseR* reveals peak bone-marrow remodeling at day 28. (a) Landmark UMAP projection of bone marrow mononuclear cells (BMMC) colored by broad landmark annotations, showing T/NK, monocyte/dendritic-cell (monos/DCs), B/plasmablast/plasma-cell (B/PB/PC) and other landmark regions. (b) Landmark UMAP projections colored by the fitted density contrast for day 28, month 3 and month 6 versus screening, displayed as density log2(+0.5) fold change, while controlling for cohort (NDMM or RRMM) and blocking for patient ID. (c) Sample-level embedding computed from the log2-transformed landmark density matrix: unsupervised PCA (left) and supervised pePC for time point (right), with samples colored by screening, day 28, month 3 and month 6. (d) Landmark UMAP projection colored by plsD1 scores for the day 28 versus screening density contrast, together with FGSEA plots for selected signatures. (e) Landmark UMAP projection colored by plsD2 scores for the same contrast, together with FGSEA plots for selected signatures. (f) Scatter plots of CAR construct expression versus density log2(+0.5) fold change for BMMC landmarks at day 28, month 3 and month 6 relative to screening. Landmarks are colored according to significance of the density contrast (q < 0.1).

Together, these findings illustrate the utility of *tinydenseR* in uncovering subtle, relevant cell states in real-world datasets by modeling and embedding sample-level variation in complex, longitudinal and heterogeneous datasets in a clustering-independent manner. Across preclinical and clinical settings, the framework resolved both broad shifts in cellular composition and finer transcriptional structure associated with treatment, disease history and severity, while retaining samples as the unit of inference. By combining landmark-level density modeling, supervised quantitative embedding and exploratory feature-space decomposition within a common manifold-based representation, *tinydenseR* enabled coherent interpretation of treatment- and disease-associated biology across technologies and experimental designs.

### Computational Performance and Reproducibility

Finally, we directly compared the computational performance of *tinydenseR* against two widely used workflows, *Seurat* and *miloR*, in a Posit Workbench session with 128GB of memory and 32 CPUs linked with OpenBLAS. We used the 1.5 million cell COVID PBMC atlas [26, 30, 31] described in *Seurat*’s tutorial. Analysis workflows started from raw counts and ended with quantifiable per-sample measurements – that is, for *Seurat*, percent of cells annotated as each cell type; for *miloR*, neighborhood counts; and for *tinydenseR*, percent cells annotated as each cell type, percent cells in each cluster as well as the fuzzy density matrix. We generated random splits of samples to achieve a range of total number of cells and repeated each workflow three times on each split, applying method-appropriate on-disk data management [32-34]. *tinydenseR* outperformed *Seurat* and *miloR* in processing time and *Seurat* in memory usage (Fig. 3a–b), underscoring *tinydenseR*’s efficiency in handling atlas-scale single-cell datasets. Of note, long processing times with *miloR* precluded memory usage assessment for larger numbers of cells despite the use of graph as refinement scheme [35].

**Figure 3.**
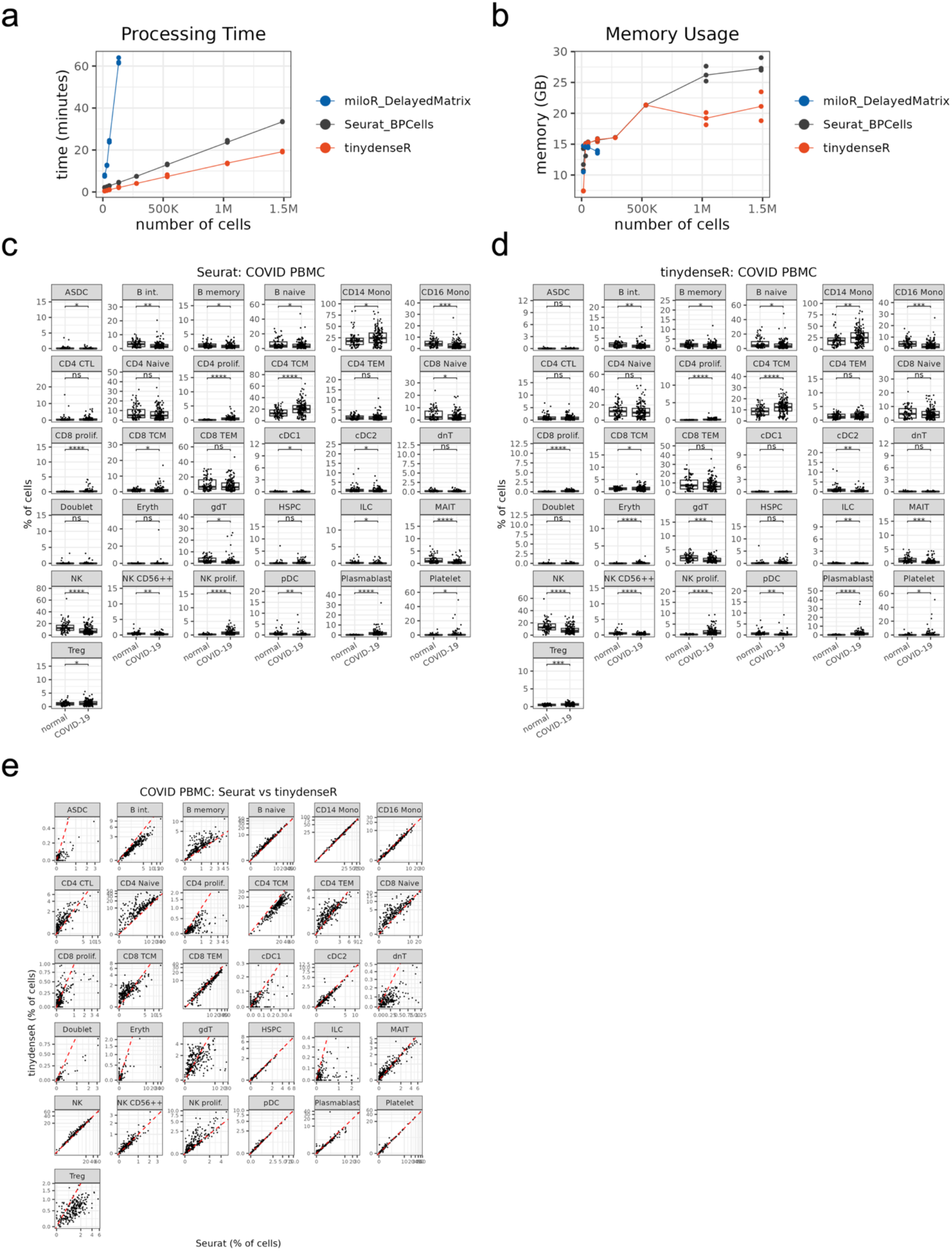
*tinydenseR* outperforms *Seurat* and *miloR* on COVID-19 PBMC multi-study atlas in speed and memory usage. (a) Processing time vs. dataset size. Wall-clock time (minutes) for three workflows, *tinydenseR, Seurat* (*BPCells* backend) and *miloR* (*DelayedMatrix* backend), across subsets of increasing number of cells from the COVID-19 PBMC atlas (up to ∼1.5 M cells). Each point represents a single run (n = 3 per method per size); lines show the median across runs. *miloR* was evaluated only for splits with fewer than ∼250k cells due to unreasonably long run times despite use of refinement_scheme = “graph”. (b) Peak memory vs. dataset size. Peak memory consumption (GB) measured during the same runs using a profvis-based peak-tracking routine. Points denote individual runs; lines show the median per method. (c–d) Percent change estimates across major immune cell subsets in COVID-19 vs. normal PBMC using Seurat (c) and *tinydenseR* (d) on the entire dataset (∼1.5M cells). Points represent individual samples. (e) Scatter plots comparing cell type abundance estimates between *tinydenseR* and *Seurat* for individual COVID PBMC samples. Red dashed line shows one-to-one correspondence between methods.

On the full dataset, similar trends were observed with *tinydenseR* and *Seurat* at the level of reference cell type annotation when comparing immune populations in healthy vs COVID PBMC (Fig. 3c–d), with some subsets showing higher correlation (e.g., cDC2, pDC, HSPC, plasmablasts, CD14 and CD16 monocytes) than others (e.g., ASDC, doublets, erythrocytes and DN T and γδ T cells) in head-to-head comparison between the methods (Fig. 3e). In part, observed differences could be due to the two workflows having different reference mapping algorithms (*symphony* [36], in *tinydenseR*, vs *Seurat*’s FindTransferAnchors and MapQuery [37]). In addition, subpopulations within CD4 and CD8 T cells are notoriously tricky to annotate at the transcriptional level, requiring protein expression profiling by CITE-seq for reliable separation[38], which could also contribute to discrepancies in cell type annotations with *tinydenseR* and *Seurat*. Of note, despite greater computational efficiency, the *tinydenseR* workflow yielded three different sample-level outputs compared to a single output in each of the other two workflows. Among *tinydenseR*’s output is the landmark-level fuzzy density matrices that can be used in linear modeling of density shifts, sample embedding and plsD decomposition to shed light on phenotype-associated variance and cell state heterogeneity. Thus, *tinydenseR* scales to ∼1.5M cells, outperforms *Seurat* and *miloR* in speed and memory, and provides richer sample-level outputs for quantitative and exploratory analysis.

## Discussion

*tinydenseR* introduces a shift in single-cell analysis by prioritizing sample-level modeling of fuzzy cell state densities around a set of shared landmarks over cell-level inference and rigid clustering. This design addresses three persistent challenges in large-scale studies: scalability, sensitivity, and robustness. By leveraging fuzzy set-based landmark representations, *tinydenseR* avoids hard cluster boundaries and preserves continuous biological variation, enabling detection of subtle heterogeneity that traditional workflows often miss.

Our results demonstrate that this approach is not only conceptually distinct but also practically impactful. Across synthetic benchmarks, *tinydenseR* accurately detected subtle cell state density shifts with high sensitivity and low estimated FDR on exhaustive label permutation tests. In preclinical xenograft models, *tinydenseR* revealed distinct transcriptional states and treatment effects between KO and WT tumors, reflecting both compositional and functional heterogeneity in response to KRAS(G12D) inhibition across genotypes. In clinical trial datasets, *tinydenseR* uncovered, for example, immune remodeling linked to disease history in multiple myeloma patients. Notably, *tinydenseR* outperformed widely used workflows such as *Seurat* and *miloR* in both speed and memory efficiency, scaling to atlas-sized datasets with about 1.5 million cells while providing richer sample-level outputs – including the fuzzy density matrix – that support reproducible and detailed biological analysis.

*tinydenseR* also provides a sample embedding that, in this work, served not only as a visualization aid but also as a mechanism for interpretation: unsupervised PCA embeddings of the landmark-by-sample fuzzy density matrix exposed dominant sample structure, including batch- and subject-level effects, whereas supervised pePC projections isolated the fitted term or contrast of interest under the same design used for inference. This supported clearer attribution of variation in the presence of batch effects, repeated-measures, and partially nested designs. By additionally reporting the percentage of total variance explained by the tested term or contrast, pePC made effect size more explicit and helped flag settings in which substantial residual variation suggested unmodeled biological or technical structure, motivating more cautious interpretation or model refinement.

This differs from sample-level representation approaches such as PILOT [39], a clustering-dependent approach which uses optimal transport to estimate distances between samples based on cell distributions over clusters, MrVI [24], which uses deep generative modeling to stratify samples and evaluate sample-level heterogeneity at single-cell resolution, and scPoli [40], which learns sample and cell representations for population-level integration and multi-scale analysis. In contrast, pePC is explicitly tied to the landmark-level fitted contrast, helping to keep it computationally lightweight, and interpretable with respect to the design matrix, while remaining complementary to *tinydenseR*’s unsupervised embedding for exposing additional technical or cohort structure relevant to interpretation [41]. Likewise, compared with metacell frameworks such as MetaCell and SEACells [20, 42], landmark-based summarization shares the broad goal of reducing data complexity for scalable downstream analysis. However, whereas metacell methods use aggregated cell groups as the primary analytic unit, *tinydenseR* uses representative landmark cells as anchors to model per-sample cell-state density distributions, enabling consistent sample-level inference across large cohorts without requiring metacell aggregates as the primary object of analysis.

For exploratory analysis of feature space, *tinydenseR* uses plsD to identify graph-smoothed expression programs that are maximally aligned with a fitted density contrast, thereby highlighting features associated with differential cell-state density due to differential abundance, differential expression, or both within the same landmark framework used for density modeling. In this respect, plsD serves a different purpose from methods such as Augur [43], which prioritize responsive cell types on the basis of perturbation separability, or TRADE [44], which estimates aggregate transcriptome-wide perturbation impact across noisy differential expression measurements. Rather than ranking cell types or summarizing transcriptome-wide effect sizes, plsD is designed to support biological interpretation and hypothesis generation without requiring clustering or cell-type labels. It does so by revealing multivariate feature programs aligned with the observed density shift through the plsD loadings and by highlighting the cell states in which those programs are most strongly represented through the plsD scores.

In parallel, *tinydenseR*’s pseudobulk DE framework provides formal sample-level inference for selected landmark subsets by aggregating expression over landmark neighborhoods using connection-strength weights derived from the same cell–landmark topology used for density estimation. This differs from conventional pseudobulk workflows that aggregate counts over predefined clusters or cell types, while still retaining samples as the unit of inference and using standard sample-level statistical modeling. In light of recent benchmarks showing that conventional pseudobulk approaches remain strong performers for differential expression analysis in many single-cell settings [12, 45], we view this component of *tinydenseR* as a way to couple inference to a manifold-informed, clustering-independent representation. In this way, the resulting DE analysis remains coherent with the upstream density model and complementary to the exploratory, subsetting-free plsD analysis.

Although *tinydenseR* is designed to be broadly applicable across technologies and biological settings, its performance still depends on well-controlled experimental design, adequate replication, and careful interpretation. As with any computational framework, the biological relevance of individual discoveries requires orthogonal validation [12]. Future work will focus on improving landmark selection, extending the framework to multimodal and spatial assays, and enabling joint analyses across technologies. Taken together, our results position *tinydenseR* as a clustering-independent, sample-centric framework for large-scale single-cell studies; a common landmark-based fuzzy density representation supports differential cell state density modeling, contrast-aligned embedding, exploratory feature interpretation, and formal sample-level inference within the same analytical scaffold. By treating samples – not cells or clusters – as the primary unit of inference, *tinydenseR* provides a coherent and practical foundation for studying complex biological variation in clinical, preclinical, and translational settings.

## Materials and Methods

### tinydenseR Algorithm

#### Step 1: Input Data

*tinydenseR* operates on quality-controlled input data. The following formats are accepted: *Seurat* (v4 and v5; on-disk or in memory), *SingleCellExperiment* (on-disk or in memory), *H5AnnData* (as h5ad on-disk file location), *BPCells, dgCMatrix, DelayedMatrix* (internally converted to *BPCells* to maximize performance) and *flowSet*/*cytoset* (cytometry only). In addition, *tinydenseR* also accepts a list of on-disk RDS file locations, one matrix for each sample in the following formats: for scRNA-seq experiments, sparse raw count matrices with genes as rows and cells as columns; and for cytometry experiments, normalized expression matrices (e.g., arcsine-transformed), with cells as rows and proteins as columns. Reproducible scripts demonstrating code usage are available in the package vignettes (link in Code Availability).

#### Step 2: Landmark Selection

The full cell-by-gene matrix for the entire dataset is never loaded into memory, unless the data input is already in memory (that is, in-memory *Seurat* or *SingleCellExperiment* objects or a stand-alone *dgCMatrix*). Regardless of data input format, *tinydenseR* processes samples individually, selecting a subset of landmark cells per sample and retaining only landmark-level data. By default, the per-sample contribution is capped at 10% of cells (configurable at object initialization) and the total number of landmarks across the dataset is capped at 5,000. Landmarks are selected with probability proportional to leverage score, where per-cell leverage score is the row-wise sum of squared left singular vectors after single value decomposition (SVD) on the (preprocessed) expression matrix[46]. A fixed random seed (default = 123) ensures reproducibility in sampling.

Our rationale for landmark caps and interaction between per-sample and global limits is the following. We choose landmark sampling to be approximately proportional to per-sample cell yield while bounding total computation. Specifically, given per-sample cell counts *n*_*s*_ and a target sampling proportion *p* (default *p* = 0.1), we set a global landmark budget *target* = *min* (∑_*s*_*n*_*s*_ *p*, 5000) and allocate *n*_*perSample,s*_ = *min* (⌈*n*_*s*_ *p*⌉, ⌈*target*/*S*⌉), where *S* is the number of samples. This construction ensures that when the global cap is binding, no single high-yield sample can dominate the landmark set: each sample has the same maximum “budget” ⌈*target*/*S*⌉, while smaller samples remain limited by ⌈*n*_*s*_*p*⌉. As a result, the total number of landmarks may be less than the global budget when many samples are small, but the representation remains controlled and transparent; users can inspect *n*_*perSample,s*_ (reported in metadata) and adjust *p* to match the study’s replicate structure and heterogeneity. The key implication is that, when the global cap binds, we do not “redistribute” landmarks from large samples to small samples in an optimization sense. Instead, we impose a uniform per-sample upper bound ⌈*target*/*S*⌉ so that no sample can contribute more than an equal share of the global “budget”, while small samples still contribute only up to ⌈*n*_*s*_*p*⌉. This allocation rule ensures bounded computation while limiting over-representation of high-yield samples in the landmark set. To flag designs where proportional landmarking may under-represent very small samples, *tinydenseR* warns when sample cell yields vary by >10x and records *n*_*perSample,s*_ for inspection.

For scRNA-seq, we first apply library size normalization per cell and log-transform counts as

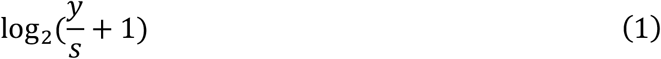

where *y* denotes raw counts and *s* is the cell-specific size factor defined as the total counts per cell divided by the mean total counts across cells. We then select highly variable genes (HVG; default = 5,000) by variance ranking, excluding by default TCR/Ig, mitochondrial, and ribosomal protein genes (approximate regex); a user-specified force-in list can be optionally included in the HVG set. We compute a truncated SVD (default *k* = 30) via *irlba* [47] on the per-sample centered and scaled HVG matrix, and sample landmarks are selected with probability proportional to the sum of squared entries of the corresponding rows of the left singular vectors.

For cytometry data, we use protein markers (defaults to all markers, but the list of markers used can be defined by the user) and apply per-marker centering and scaling (z-scoring) prior to SVD. We then compute a full-rank SVD (no truncation) and pick landmarks with probability proportional to the row-wise sum of squared left singular vectors. No HVG selection is performed for cytometry.

After per-sample landmarking, we pool the provisional landmarks and fit a dataset-wide PCA (RNA: truncated SVD on z-scored HVGs; cytometry: SVD on z-scored markers). We then compute dataset-wide leverage for every cell by streaming over samples, so the full cell-by-gene matrix is never materialized if the data input format was on-disk. For RNA, each sample is library-size normalized and log-transformed, with an additional scalar adjustment that matches the landmark mean size factor to avoid scale drift, and restricted to the learned HVG set. The sample-specific leverage score derived from the dataset-wide SVD for cell *i* is 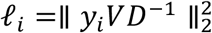, where *y*_*i*_ is the centered and scaled expression of cell *i*, and *V,D* are the dataset-wide right singular vectors and singular values, respectively.

When batch covariates are provided, we first run *harmony* [48] on the provisional landmark embedding and build a *symphony* reference. During the second landmarking pass, we project each cell with *symphony* into the *harmony*-corrected space and obtain approximate loadings by taking the SVD of the cross-covariance between the centered corrected embedding and the standardized expression. The sample-specific leverage score derived from the dataset-wide SVD after batch correction for cell *i* is then computed as above using these approximated loadings.

Using these dataset-wide leverage scores, we perform a second (final) within-sample resampling with probabilities proportional to leverage under the same per-sample and global caps, and recompute the PCA on the updated landmark set (RNA: truncated SVD with HVG re-selection; cytometry: SVD on z-scored markers). After this final PCA, if batch covariates are provided, we again run *harmony* to replace the embedding with the *harmony*-corrected one and rebuild the *symphony* reference. We then derive rotations aligned to the corrected space by linearly residualizing landmark expression against the batch design matrix and performing an SVD of the cross-covariance between the corrected embedding and the residualized, standardized expression (with sign alignment). Since *harmony* applies a nonlinear correction that alters the geometry of the original PCA space, these approximate rotations should not be interpreted as true loadings. Instead, they are intended only for exploratory analysis and visualization, not for inference on gene-PC relationships. Furthermore, while the landmark selection method described here is heuristic and performs well empirically, it does not guarantee mathematical optimality. Such guarantees would require computing exact leverage scores over the entire expression matrix, which in principle is possible but would entail prohibitive memory and I/O costs.

#### Step 3: Landmark Graph Building

Landmark expression data (after PCA/*harmony* for scRNA-seq, or SVD/harmony for batch-corrected cytometry, as described in the previous step; for cytometry without batch correction, internally z-scored by uwot::umap using scale = TRUE) is used to construct an approximate k-nearest neighbor (k-NN) graph (default *k* = 20) on Euclidean distance via the annoy algorithm and to train a UMAP model, both using the *uwot* package [49].

From this k-NN graph, we derive a Jaccard index-based Shared Nearest Neighbor (SNN) graph for clustering. Let *N*(*i*) and *N*(*j*) denote the respective sets of k-nearest neighbors of two landmarks *i* and *j*. The Jaccard similarity is:

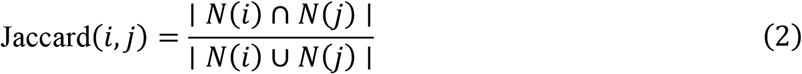

This measure captures the overlap in neighborhoods rather than raw distance, making the graph more robust to local density variations. The resulting SNN matrix is sparse and symmetric, with entries pruned below a threshold (default *τ* = 1/15) to reduce noise.

We cluster landmarks using the Leiden algorithm [50, 51] on the SNN graph, optimizing the Constant Potts Model (CPM) objective for modularity and connectivity. Initialization uses k-means (*k* = min(25, ⌈*n*/2⌉) on the Laplacian Eigenmaps (LE) embedding (described below). Clusters smaller than 0.5% of landmarks are considered stragglers and merged into the cluster with the highest mean connectivity to the stragglers. Optionally, the user can manually provide a list mapping one or more clusters to cell types. This is particularly useful in cytometry experiments, where there typically is a larger number of cells and lineage marker-based hierarchical analysis is common.

For LE graph construction, let *G* = (*V, E*) be the k-NN landmark graph described above with ∣ *V* ∣= *n* landmarks. From the directed, kNN-derived adjacency matrix *A*, we form a symmetric, unweighted adjacency

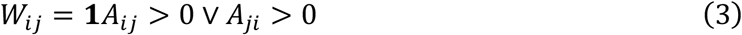

and degree matrix *D* = *diag*(*d*_1_, … , *d*_*n*_) with *d*_*i*_ = ∑_*j*_*W*_*ij*_.

We use the symmetric normalized Laplacian:

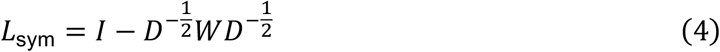

For LE embedding, we compute the smallest *n* − 1 eigenpairs of *L*_sym_ (or as many as numerically feasible), sort eigenvalues ascending 0 = *λ*_0_ ≤ *λ*_1_ ≤ ⋯, and discard near-zero eigenvalues *λ*_*ℓ*_ ≤ *ε* (with *ε* = 10^−6^ in our implementation) to remove trivial/degenerate modes.

Let the retained non-trivial eigenpairs be 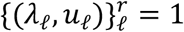, where each *u*_*ℓ*_ ∈ ℝ^*n*^. We select the embedding dimension *k* via a smooth “elbow” criterion (second derivative) on {*λ*_1_, … , *λ*_*r*_}, bounded by a target dimension *k*_⋆_ (the number of available PCs) and by *r*, and floored at 2:

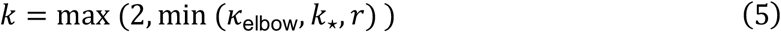

The LE coordinates are the first *k* non-trivial eigenvectors, rescaled by *D*^−1/2^ and column-normalized:

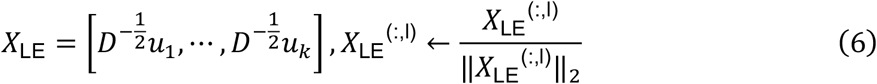

#### Step 4: Landmark Connectivity, Density Matrix Construction and Label Transfer

*tinydenseR* estimates the probability, in the fuzzy graph sense, not a calibrated probability used for statistical modeling, of each cell to be in the neighborhood of a landmark using the first part of the UMAP algorithm before dimensionality reduction. Briefly, landmarks are used as reference to query the nearest landmarks for each cell. UMAP computes a high dimensional graph with fuzzy edge weights between cells, which can be interpreted as affinities or connection strengths [49].

For each cell *c* and landmark *ℓ*, let ***z***_***c***_, ***z***_*ℓ*_ ∈ ℝ^*p*^ be the coordinates used by uwot::umap_transform:

- Cytometry case (no Harmony/Symphony): marker intensities are passed to UMAP, which internally z-scores features using center and scale from the training set (scale = TRUE) before constructing the fuzzy graph. No PCA embedding or HVG selection is applied.
- RNA case (no Harmony/Symphony): counts are library-normalized to the landmark mean library size and log-transformed, then projected by the landmark PCA:

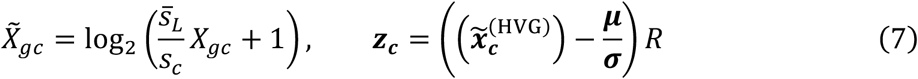

where *s*_*c*_ = ∑_*g*_ *X*_*gc*_ (cell’s library size), 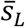 is the mean landmark library size, and (***μ, σ***, *R*) are the PCA center, scale, and rotation from the landmarks. The scalar 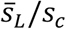 ensures per-cell normalization and aligns sample scale to the landmark mean library size.
- Harmony/Symphony case (RNA or cytometry, if batch covariates provided):

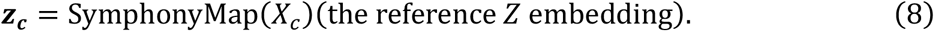

Distances used for neighbor search are Euclidean:

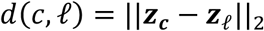

Let *N*_*k*_(*c*) ⊆ *L* be the *k* nearest landmarks of cell *c*. Define the local connectivity 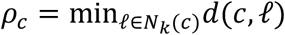. Choose *σ*_*c*_ > 0 so that the total membership mass to the *k* neighbors equals a target *τ* (UMAP/uwot default *τ* = log_2_*k*):

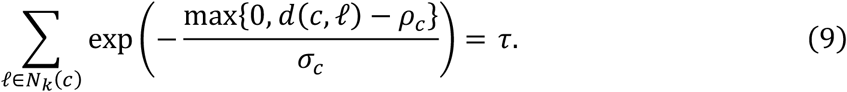

Then the directed fuzzy membership (cell → landmark) is

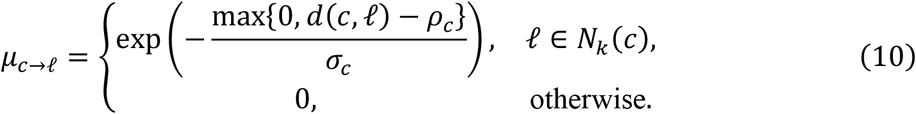

Let *μ*_*ℓ*→*c*_ be defined analogously (landmark → cell). The undirected cell-landmark fuzzy edge weight used by *tinydenseR* is the UMAP fuzzy union:

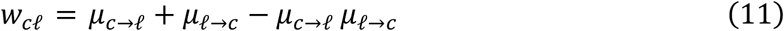

The per-cell memberships ∑_*ℓ*∈*Nk*(*c*)_*μc* → *ℓ* equal *τ* by construction.

With regard to the interpretation of the UMAP fuzzy weights as the connection strength between two points in a graph, the directed neighborhood memberships *μ*(*x, y*) are constructed by locally scaling distances so that each point distributes a fixed total membership mass across its *k* nearest neighbors. The symmetrized edge weight *P*_*xy*_ = *μ*(*x, y*) + *μ*(*y, x*) − *μ*(*x, y*)*μ*(*y, x*) corresponds to the probabilistic t-conorm (fuzzy union), i.e., the probability that at least one directed connection exists under an independence approximation. Further, treating *P*_*xy*_ as a connection strength in the high-dimensional fuzzy graph, we sum these weights from cells to landmark nodes to obtain a continuous proxy for abundance around each landmark (more on this topic below). Because the local membership mass is constrained per cell and we additionally apply size normalization across samples, the resulting landmark-by-sample matrix captures relative enrichment/depletion of cells near landmarks while remaining comparable across samples.

So, in *tinydenseR*, we sum the connection strengths for all cells in a sample to each landmark to yield a measure of sample-level abundance near each landmark. We normalize these abundances by size factors, which are the number of cells in each sample divided by the mean number of cells across samples, preserving the scale of the original data and facilitating log transformation for linear modeling. Thus, for each landmark, the sum of affinities or connection strengths across all cells in a sample is used as an estimate of the abundance of cells from that sample around that landmark.

If *w*_*cℓ*_ is the UMAP-derived fuzzy connection strength between cell *c* and landmark *ℓ* after fuzzy set union symmetrization, the unnormalized abundance around landmark *ℓ* in sample *s* is:

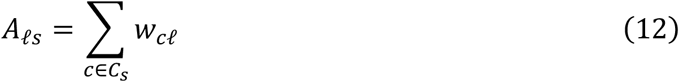

where *C*_*s*_ is the set of cells belonging to sample *s*.

To account for differences in number of cells per sample, the computed affinities are size normalized using:

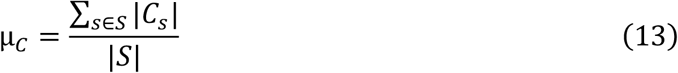

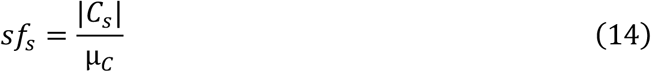

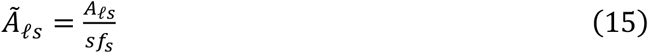

where μ_*C*_ is the across-sample mean of |*C*_*s*_|, *S* is the set of samples *s, sf*_*s*_ is the sample-specific size factor and Ã_*ℓs*_ is the size-normalized abundance. This density matrix represents the fuzzy distribution of cells across landmarks (rows) for each sample (columns).

Empirically, we found 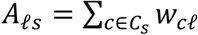 to be well-suited as abundance surrogate. Theoretically, let *w*_*cl*_∈ [0,1] denote the UMAP fuzzy-graph connection strength between cell *c* and landmark *l*. The aggregated weight

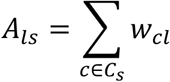

admits a soft-assignment interpretation: if *X*_*cl*_ ∈ {0,1} is a latent indicator (as a conceptual device) for whether cell *c* contributes to landmark *l*, with E[*X*_*cl*_ ∣ *z*_*c,l*_] = *w*_*cl*_ , then

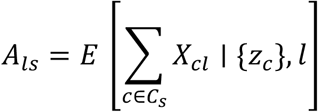

represents the expected soft assignment count of sample-*s* cells in the landmark neighborhood. This provides a theoretical justification for using *A*_*ls*_ as a continuous abundance surrogate without requiring cell independence, only the interpretation of *X*_*cl*_ as soft membership weights. UMAP’s per-cell membership-mass budget (∑_*l*∈*Nk*(*c*)_ *μ*_*c*→*l*_ ≈ *τ*) ensures bounded, locally normalized contributions per cell. Size normalization Ã_*ℓs*_= *A*_*ls*_/*sf*_*s*_ with *sf*_*s*_ ∝ |*C*_*s*_| therefore yields per-cell quantities comparable across samples, which we log-transform prior to modeling.

On the topic of sensitivity to UMAP membership construction, the fuzzy connection weights *w*_*cl*_ depend on the UMAP neighborhood construction through the choice of *k* nearest landmarks, the per-cell membership mass constraint *τ* (approximately *log*2(*k*) under *uwot* defaults), and the internally computed local scaling parameters *ρ*_*c*_ and *σ*_*c*_ that convert distances into directed memberships *μ*. We hold these choices fixed within each analysis and use the same symmetrization (fuzzy union) for all datasets. Considering how different the datasets used here are from each other, we believe the consistent results shown demonstrate robustness of the resulting landmark-by-sample densities to parameter choice. Furthermore, because *w*_*cl*_ is a locally normalized, distance-based connectivity measure rather than a physical cell count, the density matrix should be interpreted as a relative measure of enrichment/depletion of cells near landmark neighborhoods; downstream modeling therefore uses size normalization and log transformation to stabilize scale across samples prior to inference. Users can optionally adjust the UMAP/uwot parameters *k* when applying *tinydenseR* to new datasets, and the unsupervised sample embeddings (PCA/trajectory) provide a practical diagnostic to reveal dominant sample structure that may reflect unmodeled technical or biological variation.

Finally, the Leiden clustering labels from Step 3 are transferred from landmarks to cells within each sample using confidence-thresholded voting on the UMAP-derived fuzzy cell-landmark graph. For each cell, the confidence of a given cluster label is defined as the fraction of total fuzzy connection strength attributable to landmarks carrying that label, and the label is accepted only if this confidence exceeds a user-defined threshold (default = 0.5); otherwise, the cell is assigned a low-confidence label. For scRNA-seq, a reference object can also be provided at this step for cell typing. In that case, cells are first mapped to the reference using the symphony package, and cell type labels are assigned to all cells by k-nearest-neighbor voting in reference embedding space (k = 10), again subject to the same confidence threshold. That means, all cells are mapped directly against the reference, without landmarks as intermediates, and landmark labels are then updated from the subset of labeled landmark cells. For the current manuscript, a commercial PBMC reference map from Genevestigator was used to annotate leukapheresis, final product and PBMC data, and a *symphony* reference object created from the *SeuratData bmcite* CITE-seq dataset [52] was used to annotate BMMC data.

#### Step 5: Differential Density Analysis

We test for differential density at the landmark level using the fuzzy density matrix described above. Let Ã_*ℓs*_ denote the size-normalized abundance for landmark *ℓ* in sample *s*. We fit linear models with *limma* [53, 54] on log-transformed abundances,

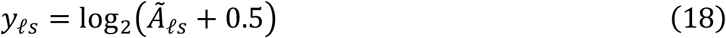

where the pseudo-count 0.5 stabilizes variance at low counts. The design matrix encodes the phenotype(s) of interest and any covariates. When repeated-measures (or other blocking) are present, we estimate a consensus intra-block correlation with duplicateCorrelation and fit blocked models with empirical Bayes moderation (eBayes, robust mode [54]).

To increase power while preserving false discovery rate (FDR) control, we implement FDR regression using *swfdr* [55, 56] with principal component (PC) scores as covariates. Specifically, let *X*_PCA_ ∈ ℝ^*n*×*k*^ be the top *k* PCs computed from the expression data as in step 2. To ensure independence from the tested covariates, we first residualize these PCs against the same design used to generate the *p*-values, respecting block structure when blocking is present. We then estimate the covariate-specific null proportion

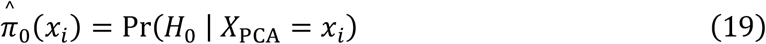

where *x*_*i*_ ∈ ℝ^*k*^, with *k* selected via a smooth “elbow” criterion (second derivative) on the PC standard deviations. The resulting 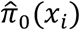 is used to rescale the *q*-values:

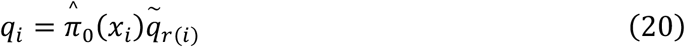

where 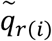 is the Benjamini-Hochberg (BH) backbone *q*-value for the rank of test *i*:

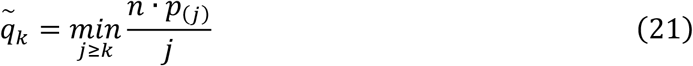

with *p*(*j*) the *j*-th ordered p-value (ascending) and *r*(*i*) the rank of *p*_*i*_ in that ordering. Since monotonicity is not enforced after scaling, these *q*-values are plug-in estimates of per-feature FDR, not a formal rejection set with guaranteed global FDR control. That means, for example, that if we reject a hypothesis with *q* = 0.1, the estimated false discovery rate for that decision is about 10% (emphasis on “about”). Unlike BH, these *q*-values do not guarantee that the set of all hypotheses with *q* ≤ *α* has FDR less or equal to alpha; they provide covariate-adjusted estimates for individual hypotheses.

In practice, for *n* < 100 tests (that is, fewer than 100 landmarks) we fall back to BH; for 100 ≤ *n* < 1000, we use standard Storey *q*-values to avoid overfitting 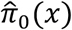; for *n* ≥ 1000, we apply *swfdr* on the residualized PCs as described above. To guard against anti-conservative behavior in regimes with large *π*_1_ (for example, strong signal, correlated tests, small *n*, etc.), we stabilize the global *π*_0_ by taking a robust median across a grid of *λ* thresholds, (that is, ranging from 0.1 to 0.8 in steps of 0.05), and if this global estimate falls below 0.6, we floor *π*μ_0_ at 0.6 before computing PCA-weighted *q*-values.

Although not used in this manuscript, *tinydenseR* also returns density-weighted BH-corrected *p*-values that down-weight landmarks in dense neighborhoods, analogous to *cydar* and *miloR*. Let *d*_*l*_ = ∑_*s*_Ã_*ℓs*_ denote the total fuzzy density mass at landmark *l*. We define weights

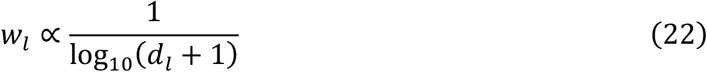

normalize them to ∑_*l*_*w*_*l*_ = 1, and apply the weighted BH procedure (equivalently, ordering by *p*_*i*_/*w*_*i*_ with the appropriate weighted step-up).

Furthermore, a “traditional” DA analysis is run based on the results from clustering (and cell typing, if provided). Here, cluster/cell type percentages are log2(+0.5)-transformed prior to fitting the same *limma* workflow models as above. This can be helpful for the analyst to get a quick overview of the results.

For differential expression, *tinydenseR* supports two pseudobulk workflows: design mode and marker mode. In design mode, pseudobulk expression is tested across experimental conditions using a user-supplied design matrix, with samples retained as the unit of inference. Cells can be restricted to a subset of interest either by specifying clustering or cell-typing labels (.id) or by providing landmark indices (.id.idx). When landmark indices are used, cells are selected via ad hoc fuzzy label transfer to the specified landmark set, rather than relying on the pre-computed cluster or cell type labels. In marker mode, two cell populations are compared within samples: group 1 is defined by .id or .id.idx, and group 2 is defined by .id2 or .id2.idx, with “..all.other.landmarks..” as the default comparator.

Pseudobulk aggregation is cell-centric in both modes. That is, cells are first selected into the population of interest, and expression from those selected cells is then aggregated using their full fuzzy connection-strength profiles across all landmarks, rather than restricting the aggregation weights to the target landmarks only. Let *I*_*s*_ denote the selected cells in sample *s*, let 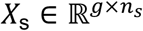 be the gene-by-cell count matrix for RNA data, and let 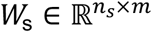 be the cell-landmark fuzzy weight matrix with entries *w*_cl_. For RNA, the pseudobulk for gene *g* in sample *s* is computed as

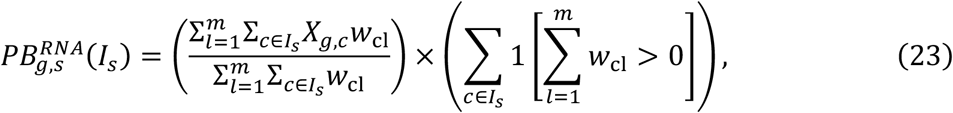

that is, a connection-strength-weighted mean multiplied by the effective number of selected cells with nonzero total fuzzy mass. For cytometry, if 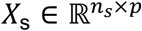 denotes the cell-by-marker matrix, the pseudobulk marker intensity for marker *p* is

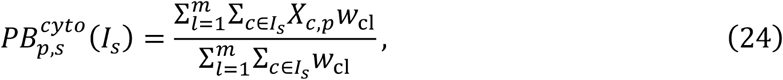

that is, a connection-strength-weighted mean marker intensity without additional cell-count scaling. In marker mode, these pseudobulks are computed independently for the two groups within each sample.

For scRNA-seq in design mode, weighted pseudobulk counts are filtered for low expression, normalized by TMM (*edgeR* [57]), and transformed with *voom* (*limma*) before fitting linear models under the user-supplied design matrix, with optional contrasts and optional blocking via limma::duplicateCorrelation; robust empirical Bayes moderation is then applied and p-values are adjusted per coefficient using BH FDR. For scRNA-seq in marker mode, pseudobulk counts are aggregated independently for each group and then combined into a single count matrix, which undergoes the same filtering, TMM normalization, and *voom* transformation before, followed by a paired *limma* model of the form ∼ .ids + .pairs, where .ids encodes the population contrast and .pairs encodes matched sample identity. Positive log fold changes indicate higher expression in group 1, whereas negative log fold changes indicate higher expression in group 2. In both modes, samples with fewer than 10% of the mean selected-cell count are excluded. Optional gene-set analysis is performed on the *voom* expression matrix using GSVA [58] and tested with the same mode-appropriate linear-model framework. For cytometry, connection-strength-weighted mean marker intensities are analyzed with the corresponding *limma* framework in design mode or marker mode, again with robust empirical Bayes moderation and FDR adjustment.

Gene sets used here that were collected from MSigDB using *msigdbr* [59] (species = “Homo sapiens”) included:

- Hallmark:
  - G2M_CHECKPOINT
  - KRAS_SIGNALING_DN
  - KRAS_SIGNALING_UP
  - PI3K_AKT_MTOR_SIGNALING
  - MTORC1_SIGNALING
  - INTERFERON_GAMMA_RESPONSE
  - INTERFERON_ALPHA_RESPONSE
- Reactome:
  - NEUTROPHIL_DEGRANULATION
- KEGG:
  - NATURAL_KILLER_CELL_MEDIATED_CYTOTOXICITY
- C8 Cell Type Signature:
  - HE_LIM_SUN_FETAL_LUNG_C2_PDC_CELL
  - HE_LIM_SUN_FETAL_LUNG_C2_HSC_ELP_CELL

Additional gene sets used here are described in Supplemental Table 1.

#### Step 6: Sample embedding (PCA/trajectory and pePC)

*tinydenseR* provides sample-level embeddings for visualization and for quantifying the amount of sample-to-sample variation attributable to specific modeled effects. All embeddings are computed from the landmark-by-sample matrix *Y* ∈ ℝ^*L*×*S*^, where *L* is the number of landmarks and *S* the number of samples; in this manuscript, *Y* corresponds to the log2-transformed, size-normalized fuzzy densities used for landmark-level modeling in Step 5.

Unsupervised PCA (sample PCA): to summarize dominant axes of sample variation without conditioning on covariates, we compute a PCA on the transposed matrix *Y*^⊤^ (samples as rows, landmarks as features) using a truncated SVD implementation (*irlba*). The PCA is centered by landmark means and not scaled. The resulting sample coordinates (scores) are stored as *PC*1, *PC*2, …, alongside landmark loadings (“rotation”), center, scale, and standard deviations. This embedding is intended as an unsupervised diagnostic of major sources of variation in landmark densities (e.g., batch structure, subject structure, or disease separation) under the same feature space used for inference.

Unsupervised diffusion-map trajectory embedding (traj): to visualize continuous sample trajectories, we compute a diffusion map on *Y*^⊤^ using destiny::DiffusionMap with user-selectable distance metric (default: cosine) and a specified number of components (*n*_*eigs*_, capped by matrix dimensions). We suppress the internal PCA step in *destiny [60]* (i.e., npcs = NA) to ensure the diffusion map is computed directly from the landmark-feature representation. Sample coordinates are constructed by scaling diffusion eigenvectors by 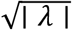, where ∣ *λ* ∣ are diffusion eigenvalues. This yields diffusion components *DC*1, *DC*2, … for sample-level trajectory visualization in the same landmark feature space.

*tinydenseR* introduces pePC, a supervised sample embedding that isolates variation attributable to a prespecified effect, computed from partial fitted values under the same linear modeling framework used for testing (Step 5). Two equivalent constructions are supported depending on how the model is specified:

##### (i) FWL-based contrast extraction (exact for OLS; approximate under blocking)

When a cell-means model is fit with contrasts (e.g., ∼ 0 + *Group* + *Batch* + *Age* with a contrast *TrtVsCtrl*, pePC is computed using the Frisch-Waugh-Lovell (FWL) decomposition to obtain the partial fitted component for a single contrast. Let *X* be the full design matrix used in limma and let *c* denote the contrast vector. We form the contrast regressor *xc* = *X*_*group*_*C*, where *Xgroup* includes the design columns participating in contrasts. We then residualize *xc* against nuisance covariates *Z* (the remaining design columns, with an intercept added if absent) to obtain *x*_*c*,⊥_. In parallel, we fit the nuisance model *Y* ∼ *Z* (optionally with a block structure via limma::duplicateCorrelation) and compute nuisance fitted values *Ŷ*_*red*_; nuisance-residualized features are *Y*_⊥_ = *Y* − *Ŷ*_*red*_. For each landmark *g*, the partial coefficient is computed as

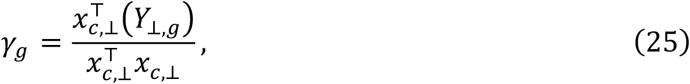

and the partial fitted matrix is the rank-1 outer product

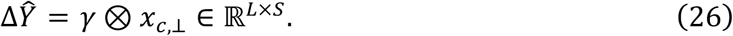

When the nuisance fit uses limma::duplicateCorrelation (blocking), the FWL decomposition is treated as an approximation to the corresponding generalized least squares decomposition.

##### (ii) Nested-model partial fitted values (difference of fitted values)

Alternatively, when models are fit without contrasts, we compute partial fitted values by comparing a full model and a strictly nested reduced model (obtained by dropping one or more terms). Denoting the fitted values under the two models as *Ŷ*_*full*_ and *Ŷ*_*red*_, the partial fitted matrix is computed directly as

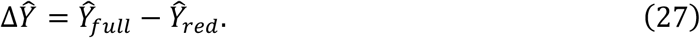

In this case, the rank of Δ*Ŷ* is bounded by the difference in model degrees of freedom and by matrix dimensions.

PCA basis from the partial fitted matrix and projection of nuisance-residualized samples: given Δ*Ŷ*, we compute a PCA on (Δ*Ŷ*)^⊤^ (samples as rows) using centering and no scaling, with PCA rank set to 1 for the contrast-based method (rank-1 by construction) and to *min* {*df*_*full*_ − *df*_*red*_, *S* − 1, *L*} for nested models. We then obtain the reported pePC coordinates by projecting nuisance-residualized samples onto the partial-effect PCA loadings:

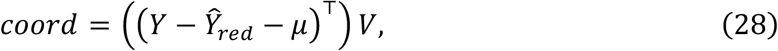

where *μ* ∈ ℝ^*L*^ is the PCA center (landmark-wise mean of Δ*Ŷ*^⊤^) and *V* are the PCA loadings learned from Δ*Ŷ*^⊤^. This yields *pePC*1, *pePC*2, … as supervised axes aligned to the specified effect, while holding nuisance terms fixed as in the reduced model.

Diagnostics and effect size summary: to assess whether nuisance-residualized sample projections align with the fitted effect direction, we report the per-component correlation between (a) PCA scores computed directly on Δ*Ŷ*^⊤^ and (b) the nuisance-residualized projection coordinates. In addition, we report the variance captured by each pePC axis as a fraction of the total variance in *Y* (computed as the sum of landmark-wise variances across samples), providing a scale-free summary of how much of the overall landmark-density variation is captured by the partial-effect axes.

#### Step 7: Graph-diffused, density contrast-aligned partial least squares decomposition (plsD)

To aid interpretation of landmark-level density contrasts, *tinydenseR* implements a graph-aware partial least squares decomposition, termed plsD. Unlike the pseudobulk DE procedure described above, plsD is not a formal hypothesis test. Instead, it is an exploratory multivariate method that identifies genes or markers whose expression patterns are aligned with a fitted density contrast across landmarks.

Let *Y*_*c*_ ∈ ℝ^*L*^ denote the centered vector of fitted coefficients for a user-specified term or contrast across the *L* landmarks obtained from the landmark-level linear model in Step 5. Let *X* ∈ ℝ^*L*×*G*^ denote the landmark-by-feature matrix, where *G* is the number of genes or markers. For scRNA-seq, features are filtered to retain genes detected in at least a user-defined minimum proportion of landmarks (defaults to 0.5%), then size-factor normalized and log2-transformed; for cytometry, landmark-level marker intensities are used (since pre-normalization is required as input to *tinydenseR* for cytometry data). Optionally, nuisance covariates may be projected out from the expression matrix before decomposition (not used in current manuscript). Let *W* denote the symmetrized shared nearest-neighbor graph on landmarks from Step 3, and let *P* be the corresponding random-walk transition matrix. Optionally, *P* may be degree-regularized by adding *τI* before row normalization, and a lazy walk may be used by replacing *P* with (1 − *α*)*I* + *αP*, where 0 < *α* ≤ 1 (not used in current manuscript).

plsD constructs a graph-smoothed local expression operator from the landmark graph and the fitted density contrast. By default, *tinydenseR* uses a density-weighted interaction of the form

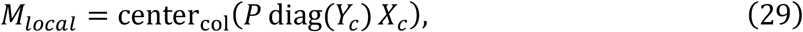

where 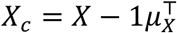 is the column-centered landmark-by-feature matrix and center_col_(·) denotes column centering of the resulting matrix. In this default mode, the density contrast enters the predictor side explicitly, so features are prioritized when their graph-smoothed expression is concentrated in landmarks with large positive or negative fitted effects. As a diagnostic alternative, the user may omit the interaction term and instead decompose

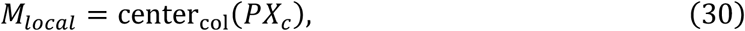

in which case the density contrast appears only on the response side of the model. This setting can also be useful when the fitted density contrast is very strong, for example when it is approximately dichotomous across landmarks or highly concentrated in a transcriptionally distinct subset of cells. Conceptually, setting .YX.interaction = FALSE removes the direct *Y*-weighting from the predictor-side operator, so plsD is driven by graph-smoothed expression patterns that covary with the contrast rather than by the signed interaction *P* diag(*Y*_*c*_) *X*_*c*_. In the current manuscript, this option was used for the synthetic flow-cytometry dataset simulating depletion of a large population (50% to 25%), for the final product versus apheresis comparison in CAR T cell manufacturing scRNA-seq data, and for the day 28 versus screening comparison in bone marrow mononuclear cell scRNA-seq data after plasma cell-targeting CAR T therapy.

plsD is implemented as a NIPALS PLS1 decomposition with implicit predictor-side deflation, so that the graph-smoothed interaction matrix is never materialized during model fitting. Instead, *tinydenseR* evaluates matrix-vector products with *M*_*local*_ and 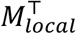 through sparse operators on the landmark graph and landmark-by-feature expression matrix, while deflating only the response vector in place. Specifically, the algorithm defines implicit operators

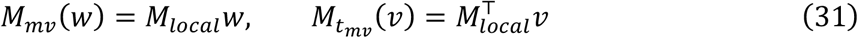

and uses these operators throughout component extraction.

Given *M*_*local*_ and *Y*, plsD applies an iterative PLS1 decomposition. At component *k*, the method estimates a unit-norm feature weight vector *w*_*k*_ ∈ ℝ^*G*^ proportional to the current predictor-response cross-product, and computes the corresponding score vector

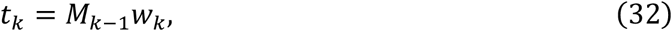

where *M*_0_ = *M*_*local*_. The corresponding predictor and response loadings are

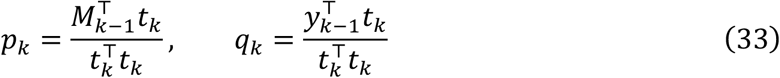

where *y*_0_ = *Y*_*c*_. Predictor-side deflation is represented implicitly through the accumulated score and loading matrices rather than by explicitly updating and storing a dense deflated matrix, whereas the response is deflated in place according to

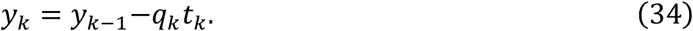

Extraction stops early if the current predictor-response cross-product is numerically negligible or if the resulting score vector has near-zero norm, in which case the returned number of components is the number successfully fitted. Components are retained in extraction order (plsD1, plsD2, …), and after fitting *tinydenseR* applies a post hoc sign convention so that positive component scores are aligned with the original density contrast. Concretely, the sign of each extracted component is set by sign{cor(*Y, t*_*k*_)}, and the same sign flip is applied to the score vectors, feature weights, predictor loadings, and response loadings.

For each fitted coefficient, *tinydenseR* stores the resulting decomposition, including coord, raw.loadings, loadings, concordance.weights, and Y.alignment. Here, coord contains the landmark-level component scores. The feature-level raw.loadings are computed post hoc as the association between each feature and the oriented component score, using either Pearson correlation, ordinary least squares regression, or Spearman rank correlation. For Pearson and OLS methods, each raw loading is then multiplied by a per-gene, per-component soft concordance.weights *c*_*jk*_ ∈ [0,1] to yield the concordance-filtered loadings. The concordance weight measures the fraction of the loading numerator 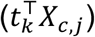 attributable to landmarks where the component score and the raw (uncentered) density-contrast coefficient agree in sign, weighted by the posterior probability that each landmark’s coefficient is genuinely nonzero and concordant: 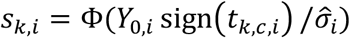, where 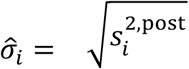 · stdev. Unscaled_*i,c*_ is the limma empirical Bayes posterior standard deviation for the coefficient of interest. Concordance weights near 1 indicate that the loading arises from landmarks with statistically confident, sign-concordant density contrasts; weights near 0 indicate that the loading is driven by structurally counterbalancing landmarks with discordant or highly uncertain coefficients. For Spearman rank correlation, concordance weights are not defined (rank-based numerators are not additively decomposable) and loadings equals raw.loadings. Positive loadings indicate features enriched along landmarks with positive component scores, whereas negative loadings indicate features enriched in the opposite direction. *tinydenseR* also reports Y.alignment, defined as

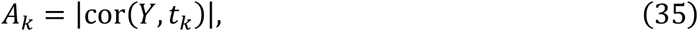

and a graph smoothness statistic for each component score vector computed from the normalized graph Laplacian.

plsD is intended for interpretation rather than formal inference. In particular, it does not produce gene-level p-values or multiple testing-adjusted significance measures. Instead, it complements pseudobulk DE by providing a graph-aware summary of the multivariate feature patterns most strongly associated with a fitted landmark-level density contrast.

### Synthetic Data

Reproducible scripts to generate the three synthetic datasets are provided in the package vignettes (link in Code Availability).

### Synthetic scRNA-seq trajectory dataset

A synthetic scRNA-seq dataset was generated with *dyntoy* using a linear trajectory with 10 milestones, 5,000 cells and 500 features (random seed = 42). Per-cell mean expression was drawn uniformly between 10 and 100, and the simulation used a differential-expression rate of 0.8. Milestone labels were obtained from the trajectory prior information. To induce differential density between two conditions (A and B), cells were assigned to condition A with milestone-specific probabilities (0.50, 0.40, 0.30, 0.20, 0.10, 0.80, 0.85, 0.90, 0.95, 1.00), with the remaining cells assigned to condition B. Thus, condition differences were introduced through a milestone-dependent redistribution of cells along the trajectory. For visualization, a principal curve was fitted to the first two PCs of the simulated log-expression matrix, and genes were reordered by Spearman correlation with the resulting trajectory coordinate before being renamed sequentially (G1-G500). Cells within each milestone were then assigned replicate labels R1-R3 in approximate proportions of 30%, 30% and the remainder, and sample labels were defined as condition-replicate combinations. The final dataset was stored as a *SingleCellExperiment* object containing counts, log-expression values, cell metadata and PCA coordinates.

### Synthetic flow cytometry differential abundance and differential expression datasets

Two synthetic flow cytometry benchmark datasets were generated to evaluate differential abundance (DA) and differential expression (DE) detection under controlled treatment and batch effects. In both simulations, samples contained approximately 50,000 cells each (sample-specific counts drawn from a normal distribution with mean 50,000 and standard deviation 500, lower-bounded at 1,000), and each cell was profiled for five markers. Samples were assigned to two treatment groups and two batches, with batch alternating deterministically across sample index. For each simulated sample, marker values were written to per-sample FCS files after log transformation.

For the DA simulation, treatment groups were Baseline and Depletion. Three baseline target-cell frequencies were considered: 0.5%, 5%, and 50%. In the Depletion group, the target-cell proportion was set to half of the corresponding baseline value (0.25%, 2.5%, and 25%, respectively). Each setting included six samples per group. Target and non-target cells were distinguished by marker-specific log-normal expression distributions for Marker1, Marker4, and Marker5, whereas Marker2 and Marker3 contributed additional cell-level heterogeneity. Batch effects were introduced as multiplicative marker-specific log-normal shifts applied to all cells in Batch2.

For the DE simulation, treatment groups were Baseline and Activation. The target-cell fraction was fixed at 5% in all samples, and DE was introduced only through Marker2 in target cells in the Activation group. Three effect sizes were simulated by shifting the log-mean of Marker2 expression in activated target cells by 0.5, 1, or 2, while keeping the log-scale standard deviation fixed at 1.5. Each effect-size setting included six samples per group. Marker1, Marker4, and Marker5 again defined target-cell identity through distinct log-normal expression distributions, and batch effects were introduced using the same marker-specific multiplicative perturbations as in the DA simulation.

### Landmark-level versus cell-level integration benchmark

To compare landmark-level and cell-level integration, we analyzed the Luecken et al. Immune_ALL_human benchmark dataset (33,506 cells × 12,303 genes; 10 batches; 16 annotated cell types) in two parallel branches. In the landmark-level branch, *tinydenseR* was run on the full count matrix using Harmony correction at the landmark level (.harmony.var = “batch”), followed by projection of all cells through the resulting Symphony reference. Because the dataset does not contain a natural sample structure distinct from batch, cells within each batch were randomly partitioned into three pseudo-samples per batch (30 pseudo-samples total), and this pseudo-sample variable was used only as sample ID in *tinydenseR*; batch correction itself was still performed using the 10 original batch labels. In the cell-level branch, all cells were normalized, log-transformed, projected by PCA, and integrated directly with Harmony using the same batch variable.

To minimize confounding from preprocessing differences, both branches used the same raw count matrix, the same random seed, the same number of principal components (30), and the same set of 5,000 highly variable genes, taken from the landmark-level branch. For the cell-level branch, PCA was computed on all cells and Harmony was run directly on the all-cell PCA embedding. For the landmark-level branch, all-cell embeddings were obtained by projecting each pseudo-sample through the landmark-level Symphony reference, reproducing the same mapping step used internally by *tinydenseR* for all-cell embedding after landmark-level integration.

Integration quality was evaluated on the shared set of cells using six metrics computed from k-nearest-neighbor graphs (k = 30): Matthews correlation coefficient (MCC) for cell-type label transfer by k-nearest-neighbor voting, average silhouette width (ASW) by cell type, inverted absolute ASW by batch (1 − |ASW|), kBET acceptance rate, graph connectivity within cell types, and batch entropy of mixing. Higher values indicate better performance for all six metrics. For visualization, UMAP was computed separately from the aligned integrated embeddings of each branch and colored by annotated cell type or batch.

### COMBAT COVID-19 PBMC scRNA-seq analysis

The COMBAT COVID-19 PBMC scRNA-seq dataset was analyzed in h5ad format using an on-disk workflow. Cells were restricted to healthy volunteers (HV) and COVID-19 samples annotated as mild, severe or critical. Quality-control filtering retained cells with at least 1,000 UMIs and 500 detected genes, and samples with at least 200 total cells. Duplicate gene symbols were removed by retaining the first occurrence. The count matrix was accessed on disk with *BPCells*, and landmark-based analysis was performed with *tinydenseR* using sample identity (`scRNASeq_sample_ID`) as the sample variable and the provided cell-type labels as reference-based annotations.

To model disease duration, time since onset of worsening symptoms was binarized into early and late strata among infected samples using the median value across non-HV samples. A design matrix was then fit with the terms ∼ 0 + Source_TSO.binary + sex + Age, where Source_TSO.binary encoded healthy, early and late disease strata within severity groups. Contrasts tested included early infection versus healthy, late versus early infection, and a severity contrasts for early critical versus early mild disease. To further interpret the early critical-versus-mild contrast, *tinydenseR* plsD was applied to the fitted density contrast. Reproducible scripts to generate the analysis results are provided in the package vignettes (link in Code Availability).

### Benchmarking using COVID PBMC atlas

PBMC datasets in h5ad format were downloaded from three cellxgene collections:

- https://cellxgene.cziscience.com/collections/8f126edf-5405-4731-8374-b5ce11f53e82
- https://cellxgene.cziscience.com/collections/b9fc3d70-5a72-4479-a046-c2cc1ab19efc
- https://cellxgene.cziscience.com/collections/03f821b4-87be-4ff4-b65a-b5fc00061da7

Preprocessing and the *Seurat* workflow followed the vignette (https://github.com/satijalab/seurat/blob/30f82df52159ac5f0feb80b149698abbd876b779/vignettes/COVID_SCTMapping.Rmd), with one additional QC step: exclusion of samples with fewer than 1,000 cells (94% of samples retained; final dataset: ∼1.48 M cells).

Scripts to reproduce the benchmarking are provided on the *tinydenseR* GitHub (see *Code Availability*). In brief:

- *tinydenseR* workflow
  - Non-default options:
    ▪ *symphony* object built from the same reference used in *Seurat* (not part of benchmarking)
    *harmony* correction by publication of origin
    2,000 HVGs (default = 5,000)
    Clustering resolution parameter set to 2 (default = 0.8, internally scaled by 1e-3 for the CPM objective)
  - All other steps used defaults: raw count normalization, transformation, dimensionality reduction, graph building, and reference mapping.
- *Seurat* workflow
  - Followed the SCT mapping vignette with *BPCells* on-disk representation for large-scale efficiency.
- *miloR* workflow
  - Non-default option:
    ▪ d = 30 in *miloR* buildGraph
    ▪ refinement_scheme = “graph”
  - Otherwise followed the *miloR* and *SingleCellExperiment* standard with *DelayedMatrix* representation:
    ▪ Normalization and log-transformation
    ▪ HVG selection via *DelayedMatrixStats* rowVars (top 2,000)
    ▪ PCA using *BiocSingular* runPCA with IrlbaParam(deferred = TRUE)
    ▪ Graph construction and neighborhood counting

Benchmark design:

- Subsets of increasing size (16 K to ∼1.5 M cells)
- Three replicates per method per size
- Metrics: wall-clock time (minutes) and peak memory (GB), measured via a

*profvis*-based peak-tracking routine [61].

### Xenograft model: SW1990 H/NRAS KO cell line engineering

Human cancer cell line SW1990 originated from the CCLE, and was authenticated by single-nucleotide polymorphism analysis and tested for Mycoplasma infection using a PCR-based detection technology (https://www.idexxbioanalytics.com/) when CCLE was established in 2012. SW1990 cells were cultured in RPMI-Glutamax Medium (Thermo Fisher Scientific) supplemented with 10% HyClone FetalClone II serum (Cytiva). Knockout clones were generated by CRISPR technology using ribonucleoproteins (RNP). CRISPR RNA (crRNA) targeting NRAS and HRAS (crRNA NRAS: AGACTCGGATGATGTACCTA, crRNA HRAS: TTGGACATCCTGGATACCGC (IDT)) were annealed with a universal tracrRNA (IDT) to form guide RNAs (gRNAs). RNPs were then formed by combining the gRNAs and Cas9 nuclease (produced in house). Cells were electroporated with the RNPs. NRAS knockout was confirmed by Western Blot. The H/NRAS double knockout was sequentially generated and verified by Western Blot but was only partial, so individual clones were isolated by plating the cells at low density in 96 well plates and selecting the wells where a single cell had formed a colony. NRAS and HRAS knockout in the single clones were further confirmed by Western blot and Amplicon-EZ sequencing (SNP/INDEL detection, Azenta) of PCR-amplified target regions.

### Xenograft model: In vivo Study and Single-Cell Dissociation

Xenograft studies were performed at Novartis in alignment with the protocols and regulations established by the Novartis Institutes for BioMedical Research Animal Care and Use Committee. The studies were approved by the Cantonal Veterinary Office of Basel Stadt, Switzerland, ensuring strict adherence to the Swiss Federal Animal Welfare Act and Ordinance. Mice used in the studies were housed under optimal hygienic conditions in individually ventilated cages, maintained on a 12-hour light/12-hour dark cycle, and provided with sterilized food and water *ad libitum*. Tumors were generated by subcutaneously injecting 6×10^6 SW1990 (WT or H/NRAS KO) cells in HBSS mixed with 50% Matrigel (BD Bioscience) into the flanks of female athymic Crl:NU(NCr)-Foxn1nu-homozygous nude mice (Charles River Germany). Tumors were measured twice a week, and tumor volume was calculated using the formula (length × width^2^) × π/6. The treatment was initiated once the average tumor size reached approximately 500 - 700 mm^3^. Mice were treated with the KRAS(G12D) inhibitor MRTX1133 at 30 mg/kg i.p. BID, formulated in 20% SBE-β-CD in 50mM citric acid. Tumors were harvested after seven days of treatment, 3h post last dose. A total of 650 mg of tumor material was cut into small pieces and processed according to the manufacturer’s protocol using the Tumor Dissociation Kit (Miltenyi Biotec), followed by mouse cell depletion (Miltenyi Biotec). Single-cell RNA sequencing libraries were prepared using the Chromium Single Cell 3’ Reagent Kit v3.1 (10x Genomics) following the manufacturer’s protocol without modifications. Briefly, single-cell suspensions were resuspended in PBS containing 0.04% BSA and quantified using the Nexcelom K2 cell counter with acridine orange/propidium iodide (AO/PI) staining to assess cell viability and concentration. Minimum cell viability was 80%. Each sample was diluted and subsequently loaded onto a Next GEM v3.1 chip (10x Genomics) to achieve a target recovery of 10,000 cells. Library quality was assessed using an Agilent 2100 Bioanalyzer. Sequencing libraries were analyzed on a NovaSeq6000 system with a depth of 50,000 reads per cell. Mapping was performed using CellRanger version 7.1.0 against the GRCh38-2020-A reference genome.

### Clinical data use statement

Data from a de-identified subset of biospecimens from the NCT04261439 (NIZ985) and NCT04318327 (PHE885) clinical trials were used solely to demonstrate computational workflows without clinical outcome analysis. No clinical endpoints, response evaluations, dose recommendations, or safety analyses were performed here. They were and/or will be described in one or more dedicated clinical manuscripts elsewhere. Sample handling and sequencing followed the study standard operating procedures as summarized below.

### NIZ985 clinical trial flow cytometry data

PBMCs were isolated from whole blood and cryopreserved until analysis. Immunophenotyping was performed starting with approximately 2×10^7^ total white blood cells (WBCs). Cells were thawed, washed and counted. Fc receptor blocking was performed, followed by viability dye staining (eFluor®506, Thermo Fisher, Cat. No. 65-0866-14). PBMCs were then stained for various surface markers, fixed, permabilized, and stained for Ki67 (clone B56, BD, Cat. No. 558615). The stained samples were analyzed on a BD LSRFortessa X-20 flow cytometer. For flow cytometry data analysis, debris was excluded based on FSC/SSC, then singlets were gated on the basis of pulse geometry, viable WBC on the basis of CD45 and viability dye, and lymphocytes were identified as SSC low. T cells were gated as CD3^+^TCRvd2^-^ in FlowJo (v10). FlowJo workspaces, including gating tree and data transformation were imported into R using *flowWorkspace* [62].

### PHE885 clinical trial scRNA-seq data

Leukapheresis and final cell product samples were frozen in 1 mL Nunc vials in CS10 by the processing facility. PBMC and BMMC were isolated from 10 mL of whole blood or bone marrow aspirates respectively and cryopreserved until analysis. For scRNA-seq analyses, a 10× Genomics Chromium instrument was used to isolate single cells from thawed samples of cryopreserved cells. Between 10,000 and 20,000 cells per sample were input into the Chromium instrument, targeting data generation between 5000 and 10,000 cells. A 10× Genomics single-cell 5′ library was constructed. The final amplified libraries were pooled, each having a unique adapter index sequence, and applied to a sequencing flow cell. The flow cell underwent cluster amplification and massively parallel sequencing by synthesis using standard approaches (Illumina Inc). Library pooling and flow cell loading were coordinated to target a median coverage of 50,000 reads per cell. The raw sequencing data were processed through 10× Genomics CellRanger, mkfastq, count, and aggr for the 5′ gene expression assay. The human reference genome is based on GRCh38. After CellRanger was run, additional quality control criteria removed droplets that met any of the three conditions: percentage of mitochondrial unique molecular identifiers >15%, number of genes detected <200, and number of unique molecular identifiers <800. For quality control, SoupX (1.4.5) [63] was used to estimate ambient RNA for each sample, and *Seurat* (5.1.0) was used for data normalization with *SCTransform* (0.4.1) [64].

## Acknowledgements

We’d like to thank Jincheng Wu, Evan Beal, Dean Lee, Noemi Di Nanni, Ethan Baker, Joshua Korn, Frank Dondelinger, Joel Wagner, Slavica Dimitrieva and Sebastian Szpakowski for critical reading of this manuscript, and Keith Kwak, Nadia Katkova, and Luz Libiran of Navigate BioPharma Services for performing peripheral blood mononuclear cell flow cytometry experiments.

## Funding

The study was sponsored and designed by Novartis Pharmaceuticals Corporation and approved by the institutional review board at each participating institution. Data were analyzed and interpreted by the sponsor and authors.

## Author contributions

Conceptualization and design of the study: PMA, SMB, AW, ML, EJO

Data acquisition: FA, SCL, MM, AW, RC, PJ, TM, NH, GC, DQ

Data interpretation: All authors

Data analysis: PMA, LHL, DS, SW

Drafting and revision of the manuscript; review of the manuscript; approval of the final version before submission: All authors.

## Competing interests

All authors report employment and stock ownership with Novartis.

## Code Availability

*tinydenseR* is available on Github (https://github.com/Novartis/tinydenseR), including code to reproduce results on synthetic and publicly available datasets as well as benchmarking.

## Data Availability

All data supporting the findings of this analysis are available within the article and its Supplementary Information. In accordance with the Health Insurance Portability and Accountability Act, Novartis does not have institutional review board approval or patient consent to share individualized patient genomic data, so data cannot be reported in a public data repository. All data provided are anonymized to respect the privacy of patients who have participated in the trial in line with applicable laws and regulations. Phase 1 studies, by their nature, present a high risk of patient re-identification; therefore, patient individual results for phase 1 studies cannot be shared. In addition, clinical data, in some cases, have been collected under contractual or consent provisions that prohibit transfer to third parties. Such restrictions may preclude granting access under these provisions. Where co-development agreements or other legal restrictions prevent companies from sharing particular data, companies will work with qualified requestors to provide summary information where possible. The data availability of these trials is according to the criteria and process described at www.clinicalstudydatarequest.com.

## Clinical trial registration information

The studies have been registered at ClinicalTrials.gov with the IDs NCT04261439 and NCT04318327.

## Supplemental Information

### R System, Packages and Versions

| *System Component* | *Value* |
| --- | --- |
| R version [65] | R version 4.5.3 (2026-03-11) |
| Platform | x86_64-pc-linux-gnu |
| Running under | Red Hat Enterprise Linux 9.6 (Plow) |
| BLAS/LAPACK | FlexiBLAS OPENBLAS |
| base packages | stats4, parallel, grid, stats, graphics, grDevices, utils, datasets, methods, base |

| *Package* | *Version* |
| --- | --- |
| Biobase [66] | 2.68.0 |
| BiocGenerics [66] | 0.54.0 |
| collapse [67] | 2.1.6 |
| dplyr [68] | 1.1.4 |
| curl [69] | 6.4.0 |
| dyntoy [70] | 0.9.9 |
| edgeR [57] | 4.6.2 |
| generics [71] | 0.1.4 |
| fgsea [72] | 1.36.0 |
| GenomeInfoDb [73] | 1.44.0 |
| GenomicRanges [74] | 1.60.0 |
| ggh4x [75] | 0.3.1 |
| ggnewscale [76] | 0.5.1 |
| ggplot2 [77] | 3.5.2 |
| ggrepel [78] | 0.9.7 |
| gridExtra [79] | 2.3 |
| GSVA [58] | 2.2.0 |
| gtable [80] | 0.3.6 |
| harmony [81] | 1.2.3 |
| igraph [82] | 2.1.4 |
| IRanges [74] | 2.42.0 |
| irlba [83] | 2.3.5.1 |
| knitr [84] | 1.5 |
| limma [53] | 3.64.1 |
| Matrix [85] | 1.7-3 |
| MatrixGenerics [86] | 1.20.0 |
| matrixStats [87] | 1.5.0 |
| patchwork [88] | 1.3.1 |
| pheatmap [89] | 1.0.13 |
| qvalue [90] | 2.40.0 |
| Rcpp [91] | 1.0.14 |
| RhpcBLASctl [92] | 0.23-42 |
| rlang [93] | 1.1.6 |
| S4Vectors [94] | 0.46.0 |
| scales [95] | 1.4.0 |
| scattermore [96] | 1.2 |
| Seqinfo [97] | 1.0.0 |
| SeuratObject [98] | 5.1.0 |
| sp [99] | 2.2-0 |
| sparseMatrixStats [100] | 1.20.0 |
| SummarizedExperiment [101] | 1.38.1 |
| swfdr [102] | 1.34.0 |
| symphony [103] | 0.1.1 |
| tidyr [104] | 1.3.1 |
| tidyselect [105] | 1.2.1 |
| uwot [106] | 0.2.3 |

**Supplemental Table 1:** Additional gene sets.

|  |  |
| --- | --- |
| plasma blast/cell | SCDC1, MZB1, IGHA1, IGHA2, IGHG1, IGHG2 IGHG3, IGHG4, IGKC, IGLC2, XBP1, PRDM1, IRF4, JCHAIN, CD38 |
| inf. mono | FCGR3A, CX3CR1, IFITM2, IFITM3, LST1, CDKN1C, S100A4, S100A6, CD14, CD1A, CD1B, CD1C, CD4, HLA-DOA, HLA-DOB, CD80, CD86, HLA-DRA, HLA-DRB1, HLA-DRB5, HLA-DRB6, CD207, CD40, CCR7, CMKLR1, ITGA4, CD83, LY75, ITGAM, ITGAX, CD209, PDCD1LG2, NRP1 |
| Tmem | NKG7, PRF1, GNLY, GZMK, GZMH, CST7, FGFBP2, CCL5, KLRG1, KLRD1, EOMES, TBX21, ZEB2, SLAMF7, KLRF1, FASLG, CD44, CCR6, CXCR3 |

## Supplemental Figures

**Figure S1.**
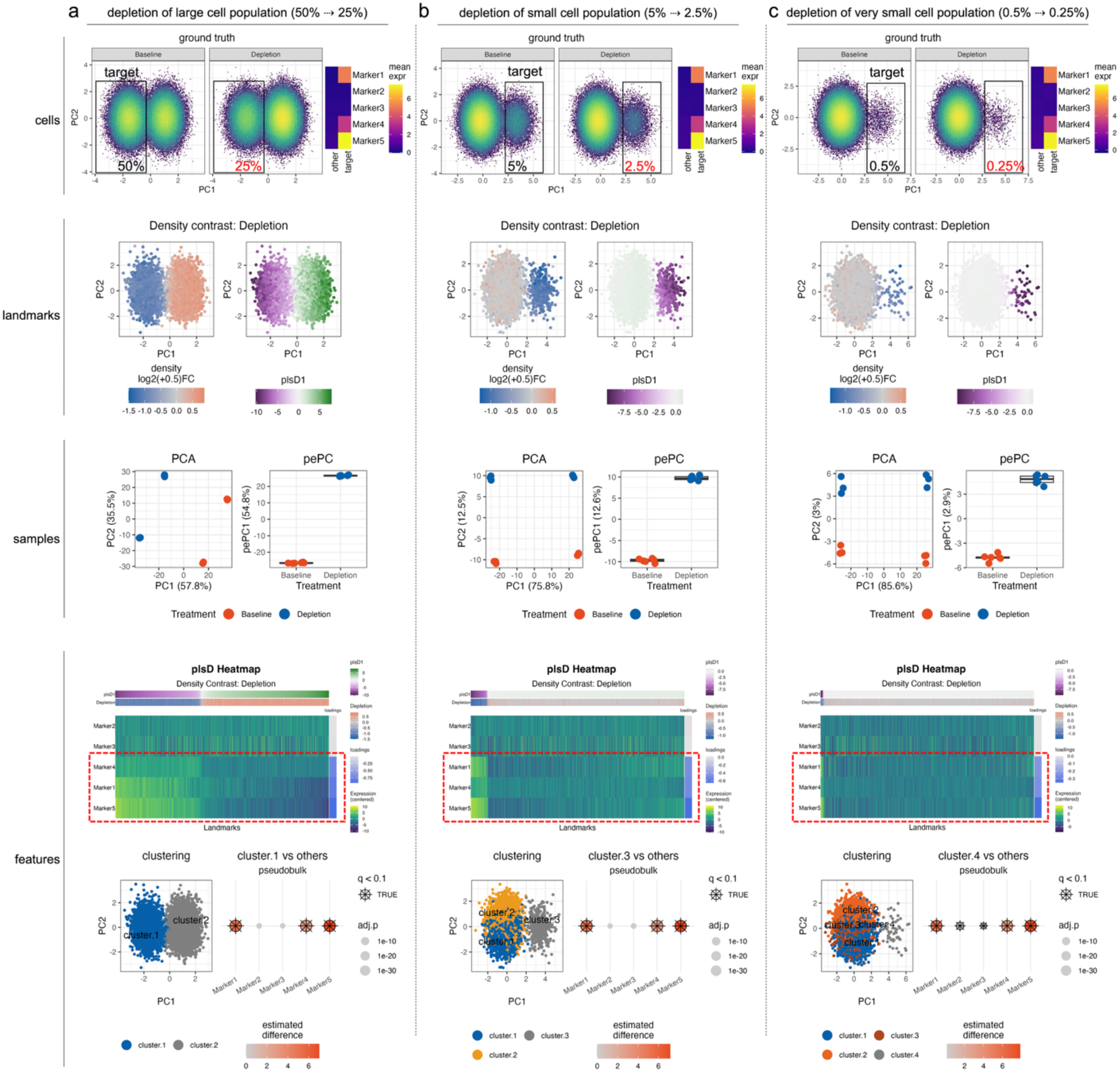
*tinydenseR* detects DA upon two-fold depletion across large, small, and very small cell populations in simulated flow cytometry data. (a-c) Simulated flow cytometry datasets with a two-fold reduction in the abundance of a target population between Baseline and Depletion. Baseline target-cell frequencies were 50% (a), 5% (b), and 0.5% (c), corresponding to frequencies of 25%, 2.5%, and 0.25% after depletion, respectively. Each scenario was simulated with five markers, two batches, and six samples per treatment group. Cells: ground-truth cell-level distributions for Baseline and Depletion, showing the target population and the imposed difference in target-cell frequency, and heatmap of mean expression levels of each marker in target vs other cells. Landmarks: log2-fold change in density around each landmark (that is, the estimated density contrast for depletion while controlling for batch; left hand side) and plsD1 scores for the same density contrast (right hand side). Samples: sample-level embeddings computed from the landmark-by-sample density matrix, shown as unsupervised PCA and supervised pePC for the depletion effect. Features: plsD heatmaps showing per-landmark plsD1 scores and density contrast as annotation strips at the top, followed by the centered-expression level for each marker (row-ranked by plsD1 loading shown as annotation strip on the right hand side). Please note that markers for the target population have negative loadings in plsD1, in accordance with the effect of depletion on the population structure. At the bottom, results of pseudobulk marker-mode DE analysis comparing clusters of target cells vs all other clusters showing expression levels of target-specific markers.

**Figure S2.**
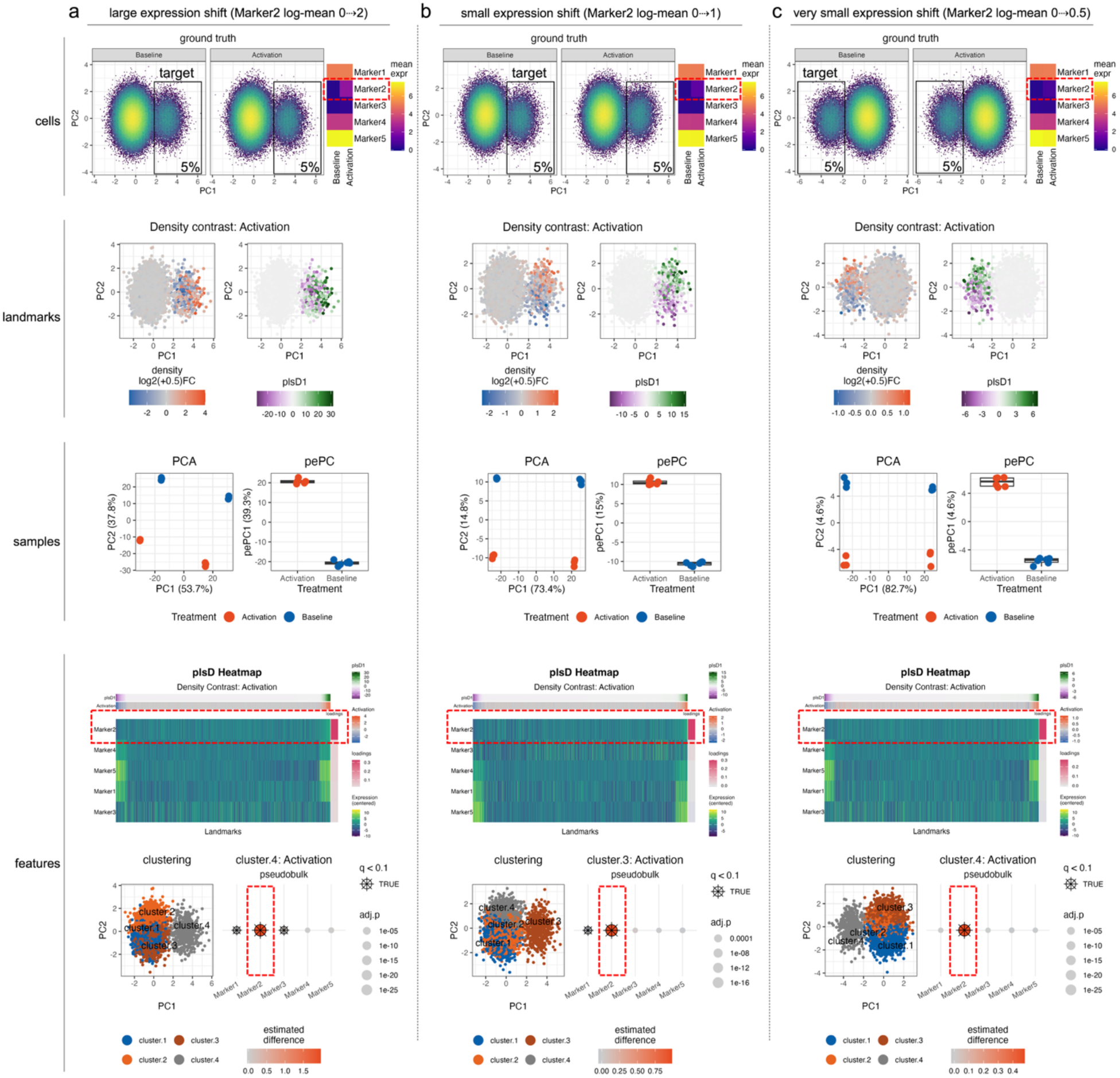
*tinydenseR* identifies DE of Marker2 upon activation in small cell populations across a range of effect sizes in simulated flow cytometry data. (a-c) Simulated flow cytometry datasets in which the target population was fixed at 5% of cells in both treatment groups and Marker2 was upregulated specifically in target cells in the Activation group relative to Baseline. Activation-induced shifts in the log-mean of Marker2 expression were 2 (a), 1 (b), and 0.5 (c), with all scenarios simulated using five markers, two batches, and six samples per treatment group. Cells: ground-truth cell-level distributions for Baseline and Activation, showing the target population and the same target-cell frequency across conditions, and heatmap of mean expression levels of each marker in target vs other cells, showing Marker2 expression increase in the target cell population with activation. Landmarks: log2-fold change in density around each landmark (that is, the estimated density contrast for activation while controlling for batch effect; left hand side) and plsD1 scores for the same density contrast (right hand side). Samples: sample-level embeddings computed from the landmark-by-sample density matrix, shown as unsupervised PCA and supervised pePC for the activation effect. Features: plsD heatmaps showing per-landmark plsD1 scores and density contrast as annotation strips at the top, followed by the centered-expression level for each marker (row-ranked by plsD1 loading shown as annotation strip on the right hand side). Please note that Marker2 has a positive loading in plsD1, in accordance with the effect of activation on the target population. At the bottom, results of pseudobulk design-mode DE analysis comparing the effect of activation vs baseline within clusters of target cells showing increase in expression levels for Marker2.

**Figure S3.**
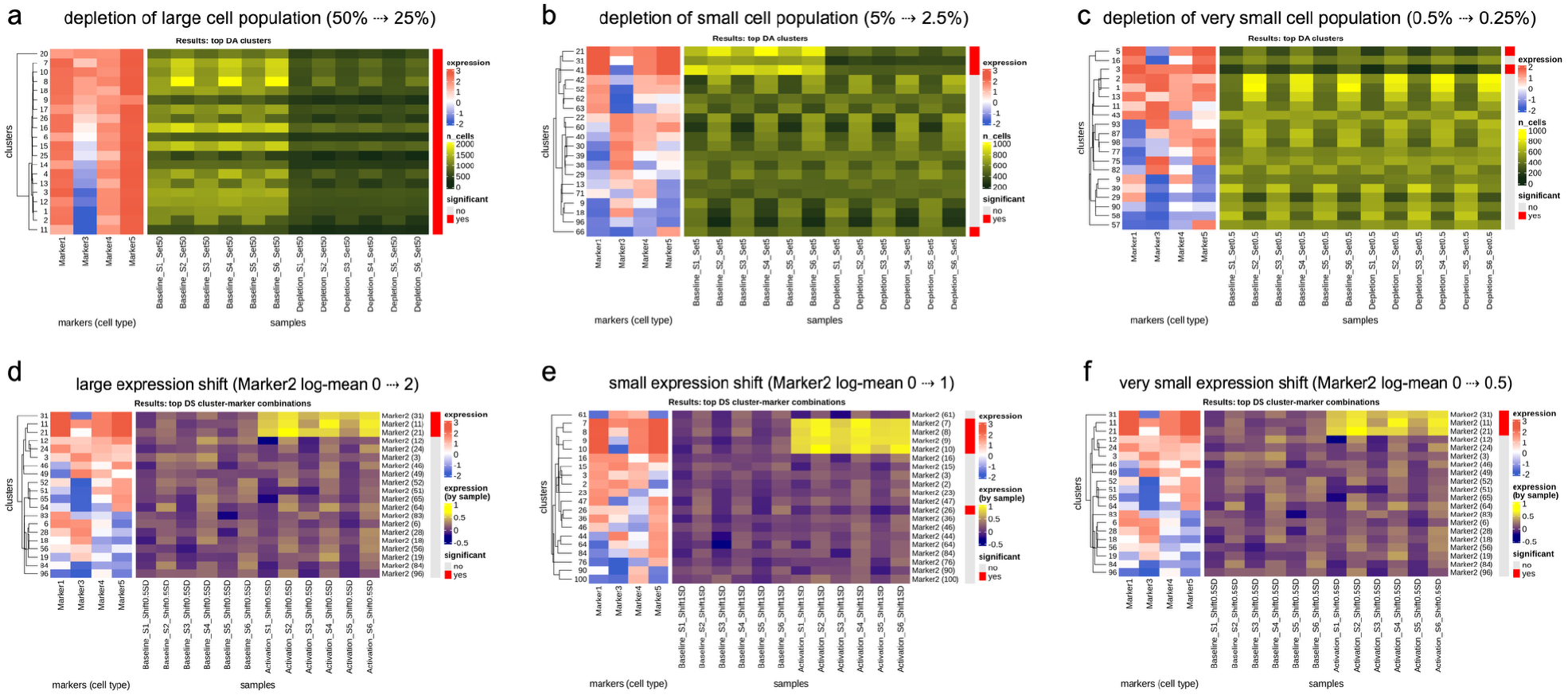
Performance of *diffcyt* on synthetic flow cytometry datasets simulating differential abundance (depletion) and differential expression (expression shift). (a–c) DA analysis: *diffcyt* results for simulated depletion of a large (50% → 25%) (a), small (5% → 2.5%) (b), and very small (0.5% → 0.25%) (c) target population. Heatmaps show the top differentially abundant clusters across samples and conditions, together with the corresponding cluster-level marker-expression profiles; significant clusters are indicated. (d–f) DE analysis: *diffcyt* results for simulated activation-dependent upregulation of Marker2 in a small target population (5% of total cells), with increasing effect sizes corresponding to shifts in the log-mean of Marker2 expression from 0 to 2 (d), 0 to 1 (e), or 0 to 0.5 (f), with log-SD fixed at 1.5. Heatmaps show the top significant cluster–marker combinations and their sample-level expression patterns, with significance indicated. In this benchmark, Marker2 was specified as the only state marker for DE testing, whereas the remaining markers were specified as type markers for clustering.

**Figure S4.**
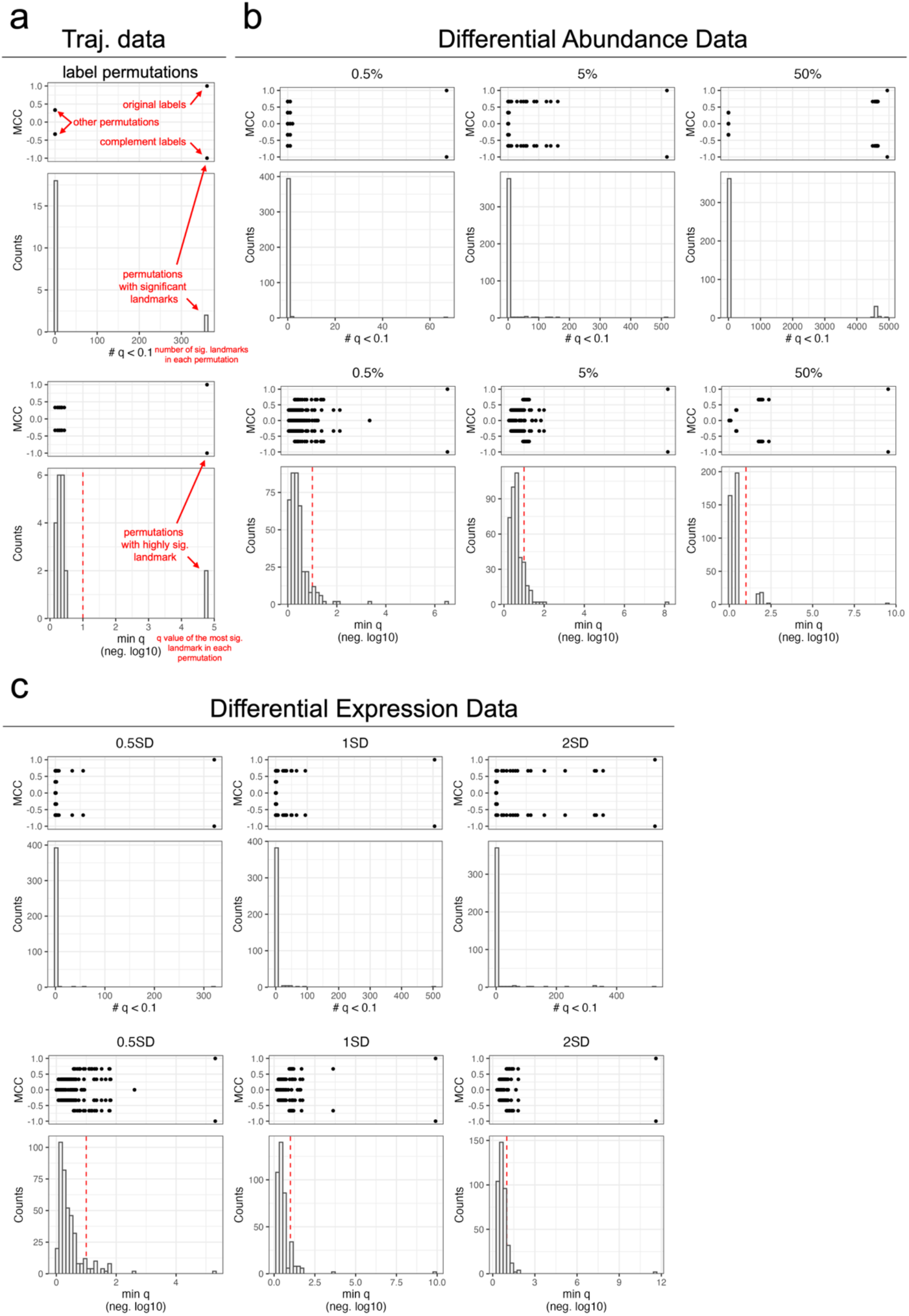
Permutation-based null-distribution discoveries across simulated scenarios. (a–c) Exhaustive label-permutation results for the synthetic trajectory (a), DA (b) and DE (c) datasets. For each condition, the upper pair of panels shows the number of discoveries (# q < 0.1) plotted against MCC (top) and its histogram (bottom), and the lower pair shows the minimum q-value on the negative log10 scale plotted against MCC (top) and its histogram (bottom). Panels correspond to the trajectory dataset (a; n = 20 permutations; m = 464 landmarks), the DA dataset at 0.5%, 5% and 50% target-cell frequencies (b; n = 400 permutations per setting; m = 5004 landmarks), and the DE dataset at 0.5SD, 1SD and 2SD effect sizes (c; n = 400 permutations per setting; m = 5004 landmarks). MCC quantifies similarity between the permuted and observed labels, with 1 denoting identical labels and -1 denoting the exact complement. Red dashed lines indicate q = 0.1 on the minimum-q histograms. Permutations were stratified within batch for the DA and DE datasets (the trajectory dataset did not have batch effect).

**Figure S5.**
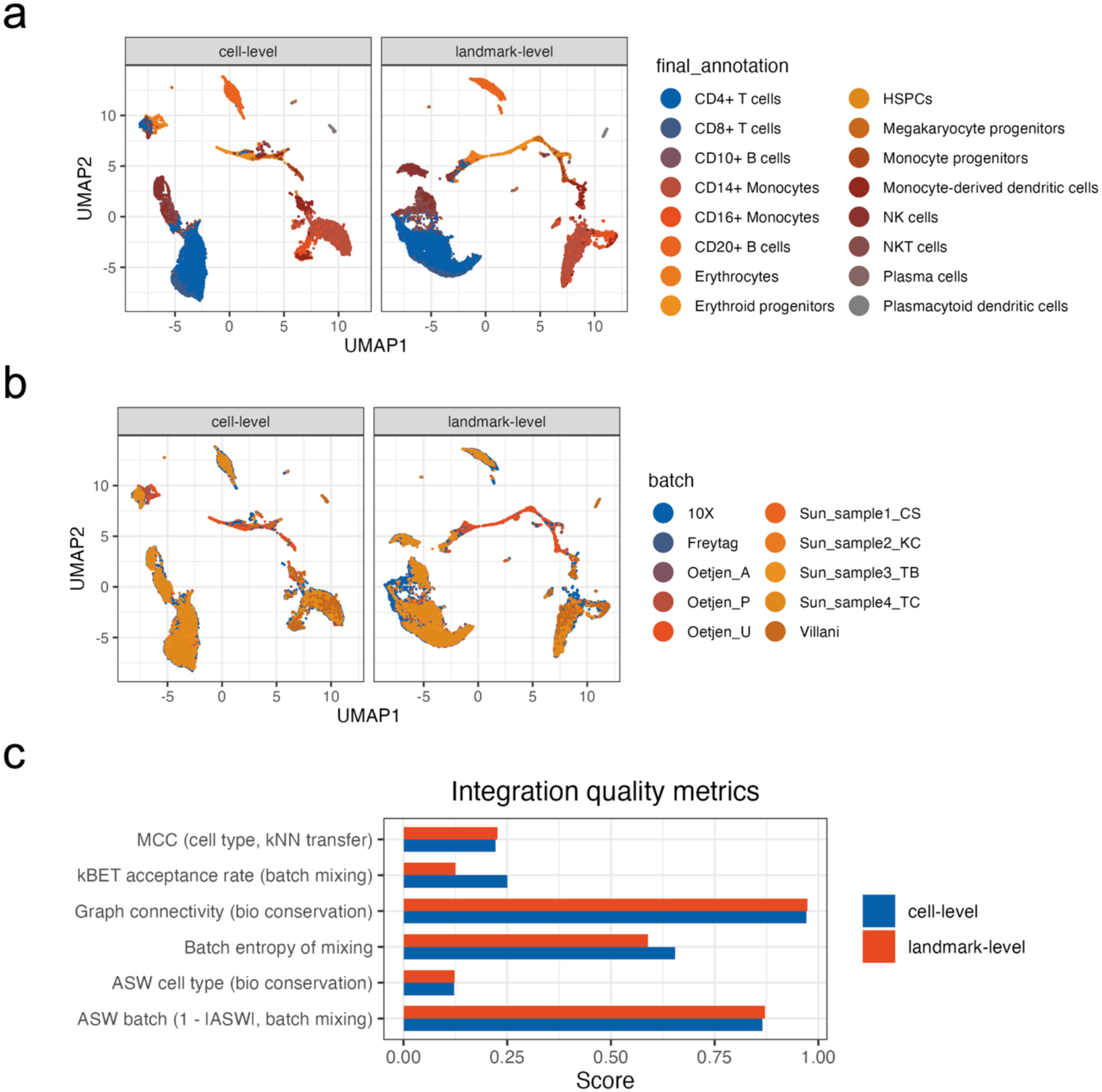
Landmark-level integration preserves major biological structure while modestly reducing local batch mixing relative to direct cell-level integration. (a) UMAP visualization of the integrated Immune_ALL_human benchmark dataset from Luecken et al. (33,506 cells; 10 batches), colored by the provided cell-type annotation (final_annotation). Left, cell-level integration obtained by Harmony on all cells. Right, landmark-level integration obtained by *tinydenseR*, in which Harmony was applied to landmark PCs and all cells were subsequently projected through the landmark-level Symphony reference. (b) The same UMAP layouts as in panel a, now colored by batch. (c) Integration-quality metrics comparing the two approaches. Biological-conservation metrics comprised Matthews correlation coefficient (MCC) for k-nearest-neighbor cell-type label transfer, average silhouette width (ASW) by cell type, and graph connectivity. Batch-mixing metrics comprised ASW by batch (1 − |*ASW*|), kBET acceptance rate, and batch entropy of mixing.

**Figure S6.**
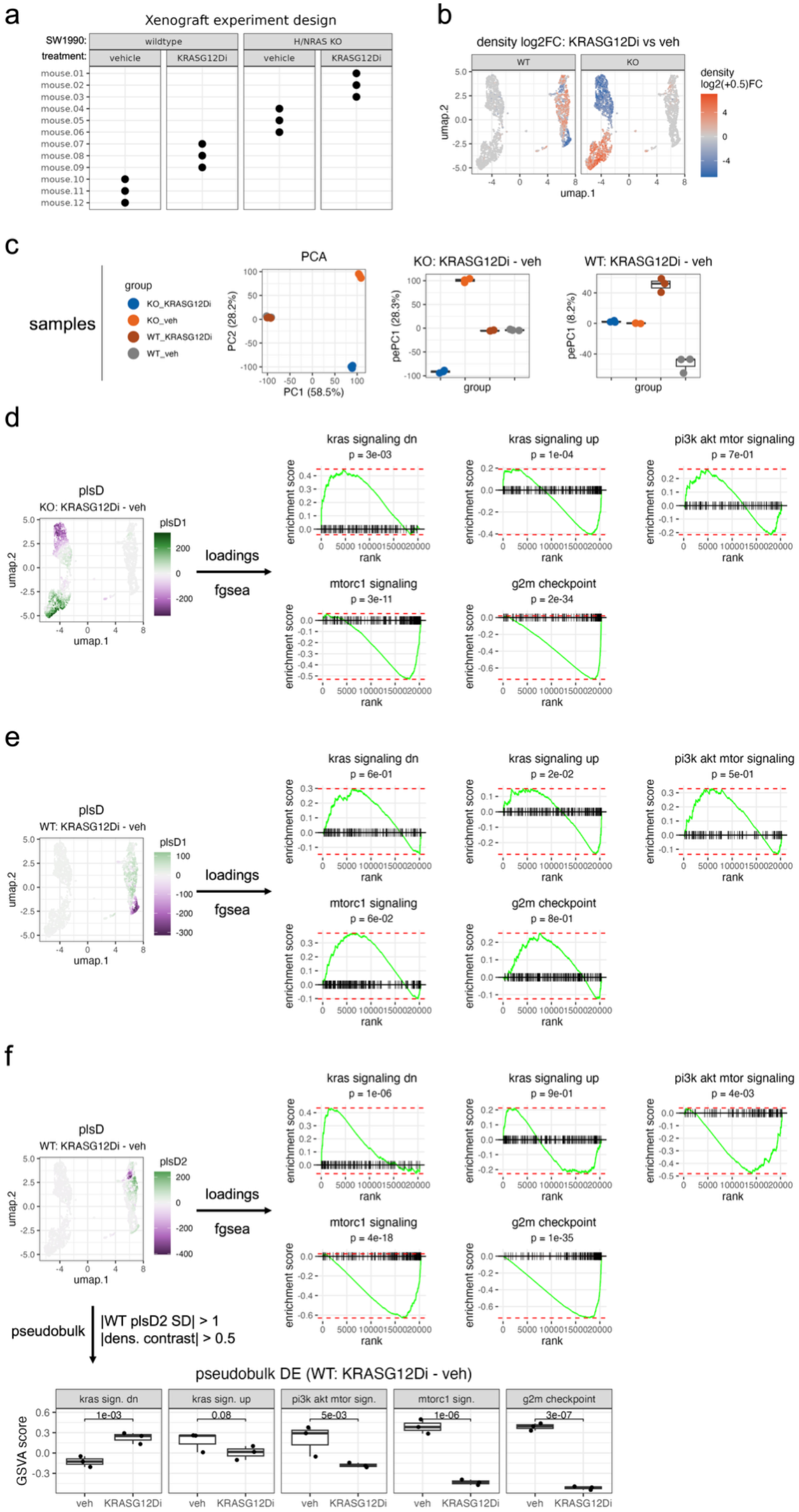
*tinydenseR* resolves genotype-specific treatment responses and WT-specific secondary transcriptional structure in a xenograft model of pancreatic cancer. (a) Experimental design: KRAS(G12D) mutant SW1990 pancreatic cancer cells, either wild-type (WT) or *HRAS*/*NRAS* double knockout (H/NRAS KO), were implanted into mice and treated with either vehicle or the KRAS(G12D) inhibitor MRTX1133 at 30 mg/kg i.p. BID for seven days. (b) Landmark UMAP projection of xenograft-derived cells colored by the fitted density contrast for KRASG12Di versus vehicle, shown separately for WT and KO tumors. Colors indicate density log2(+0.5) fold change, with positive and negative values denoting opposite sides of the treatment contrast. (c) Sample-level embedding computed from log2-transformed landmark densities: unsupervised PCA (left) and supervised pePC1 for the indicated treatment contrasts within KO and WT backgrounds (KO: KRASG12Di vs vehicle; WT: KRASG12Di vs vehicle), with samples colored by genotype/treatment group. (d) Landmark UMAP projection colored by plsD1 scores for the KO treatment contrast, together with FGSEA plots of hallmark pathways ranked by plsD1 loadings. (e) Landmark UMAP projection colored by plsD1 scores for the WT treatment contrast, together with FGSEA plots of hallmark pathways ranked by plsD1 loadings. (f) Landmark UMAP projection colored by plsD2 scores for the WT treatment contrast, together with FGSEA plots of hallmark pathways ranked by plsD2 loadings, highlighting a secondary WT-associated axis. Bottom, pseudobulk design-mode DE GSVA analysis comparing KRASG12Di vs vehicle for the subset of landmarks with |WT plsD2 score (SD)| > 1 and |WT density contrast| > 0.5 in the same direction.

**Figure S7.**
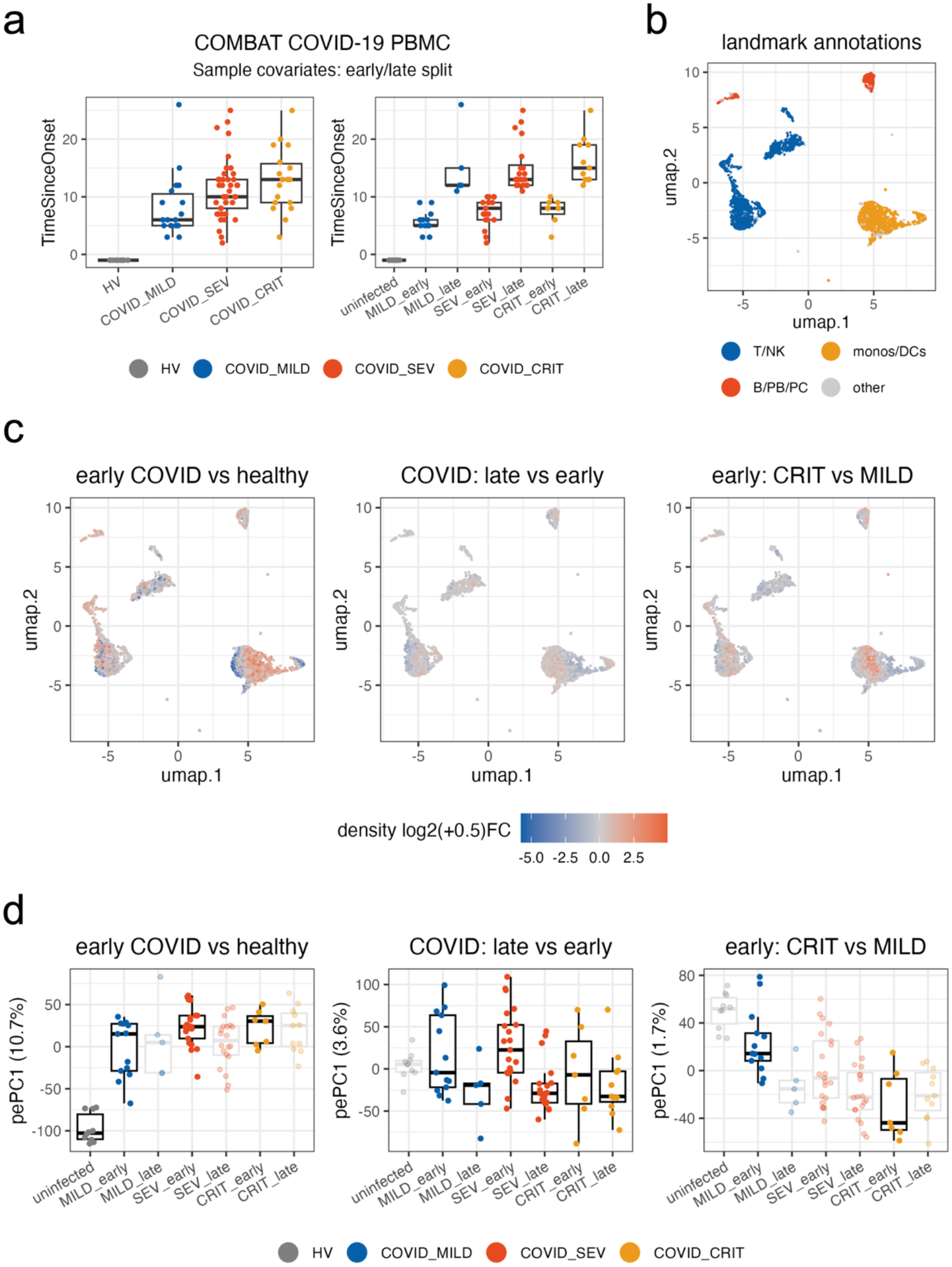
*tinydenseR* models COMBAT COVID-19 PBMC scRNA-seq data and quantitatively embeds samples along disease status, disease duration and severity. (a) Sample covariates used to define early and late disease strata. Left, distribution of time since onset of worsening symptoms across healthy volunteers (HV) and COVID-19 samples stratified by clinical severity. Right, the same samples after binarization of time since onset into early and late strata among infected samples. Points denote individual samples; boxplots summarize distributions. (b) Landmark UMAP projection colored by broad landmark annotations, showing regions enriched for T/NK cells, monocytes/dendritic cells (monos/DCs), B/plasmablast/plasma-cell populations (B/PB/PC), and other landmarks. (c) Landmark-level density contrasts for three contrasts estimated from linear models on log2-transformed fuzzy densities while controlling for sex and age: early COVID versus healthy, COVID late versus early, and early critical versus early mild disease. Landmarks are colored by density log2(+0.5) fold change. (d) Supervised quantitative sample embedding (pePC1) for the same three contrasts shown in panel (c). Samples are colored by clinical group.

**Figure S8.**
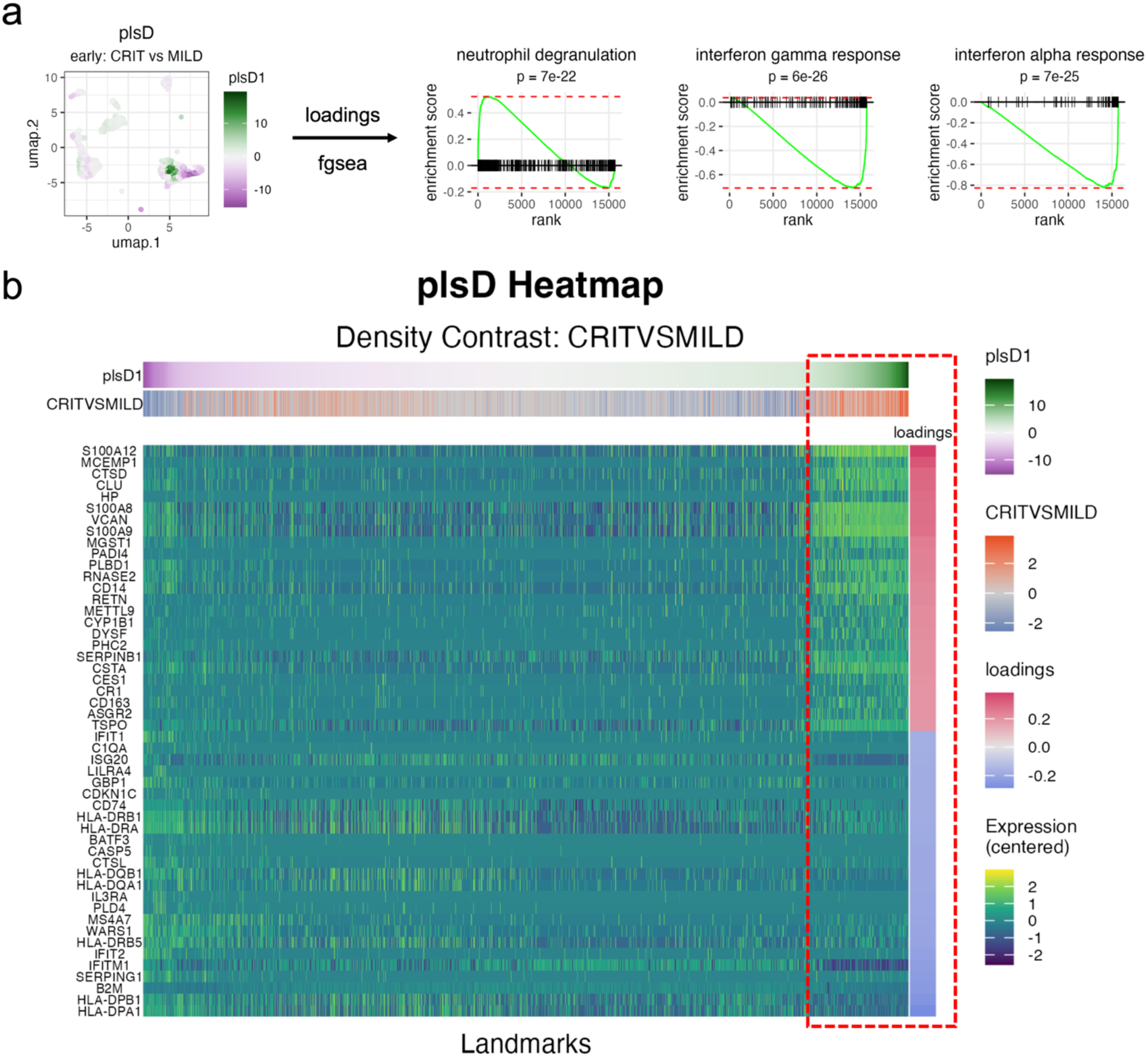
plsD1 identifies a dysregulated monocyte-associated transcriptional program in early critical versus mild COVID-19. (a) Left, landmark UMAP projection colored by plsD1 scores for the early critical-versus-mild density contrast. Right, FGSEA plots for selected pathways ranked by plsD1 loadings. (b) plsD heatmap for the same contrast, ordered by plsD1 score across landmarks. Annotation strips show plsD1 scores, the early critical-versus-mild density contrast (both at the top), and plsD1 loadings (right-hand side of the heatmap). Rows show centered expression of top 25 features associated with the component in each direction.

**Figure S9.**
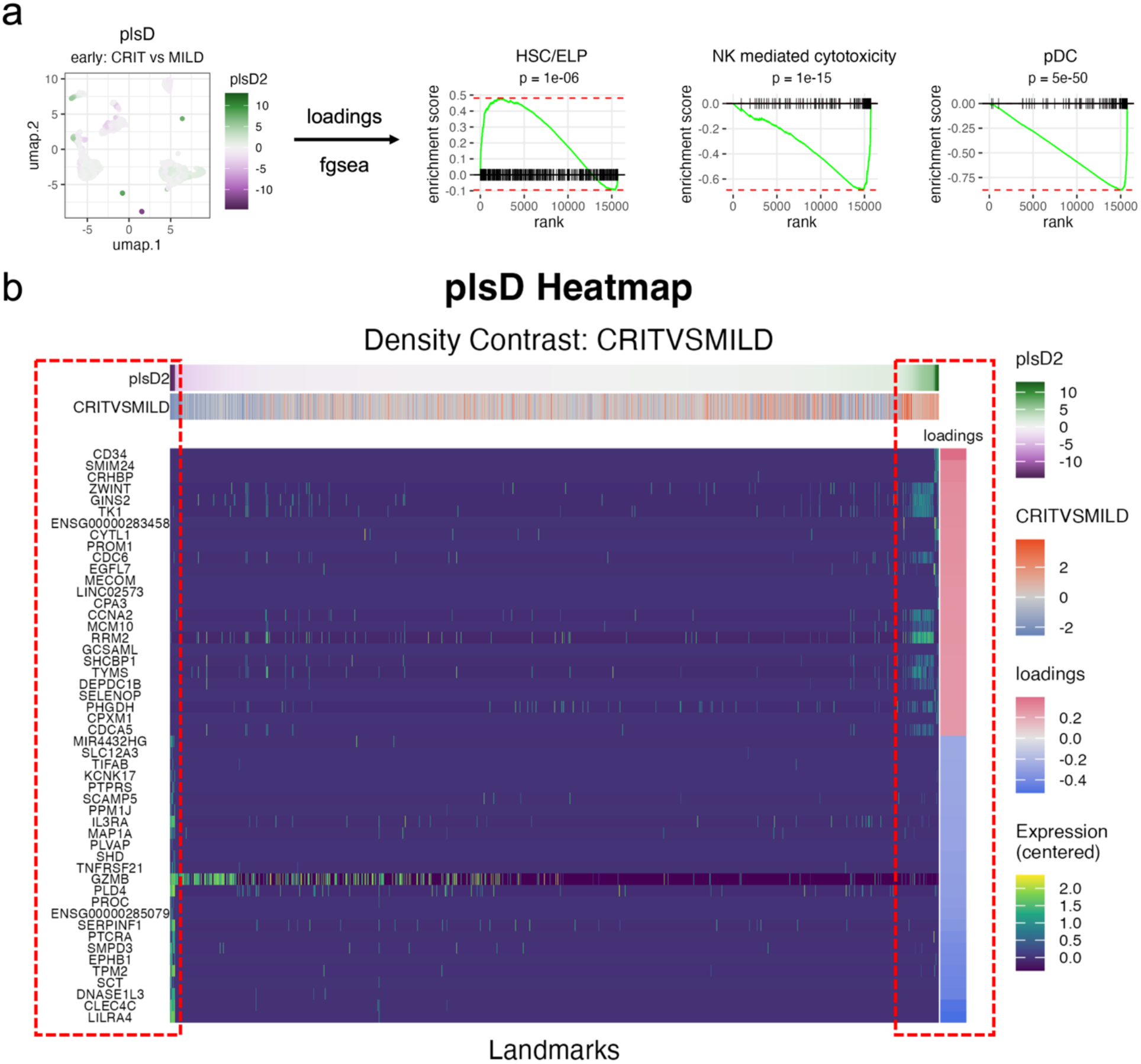
plsD2 resolves secondary transcriptional structure in the early critical-versus-mild COVID-19 contrast. (a) Left, landmark UMAP projection colored by plsD2 scores for the early critical-versus-mild density contrast. Right, FGSEA plots for selected pathways ranked by plsD2 loadings. HSC: hematopoietic stem cell; ELP: early lymphocyte progenitor; NK: natural killer; pDC: plasmacytoid dendritic cell. (b) plsD heatmap for the same contrast, ordered by plsD2 score across landmarks. Annotation strips show plsD2 scores, the early critical-versus-mild density contrast (both at the top), and plsD2 loadings (right-hand side of the heatmap). Rows show centered expression of top 25 features associated with the component in each direction.

**Figure S10.**
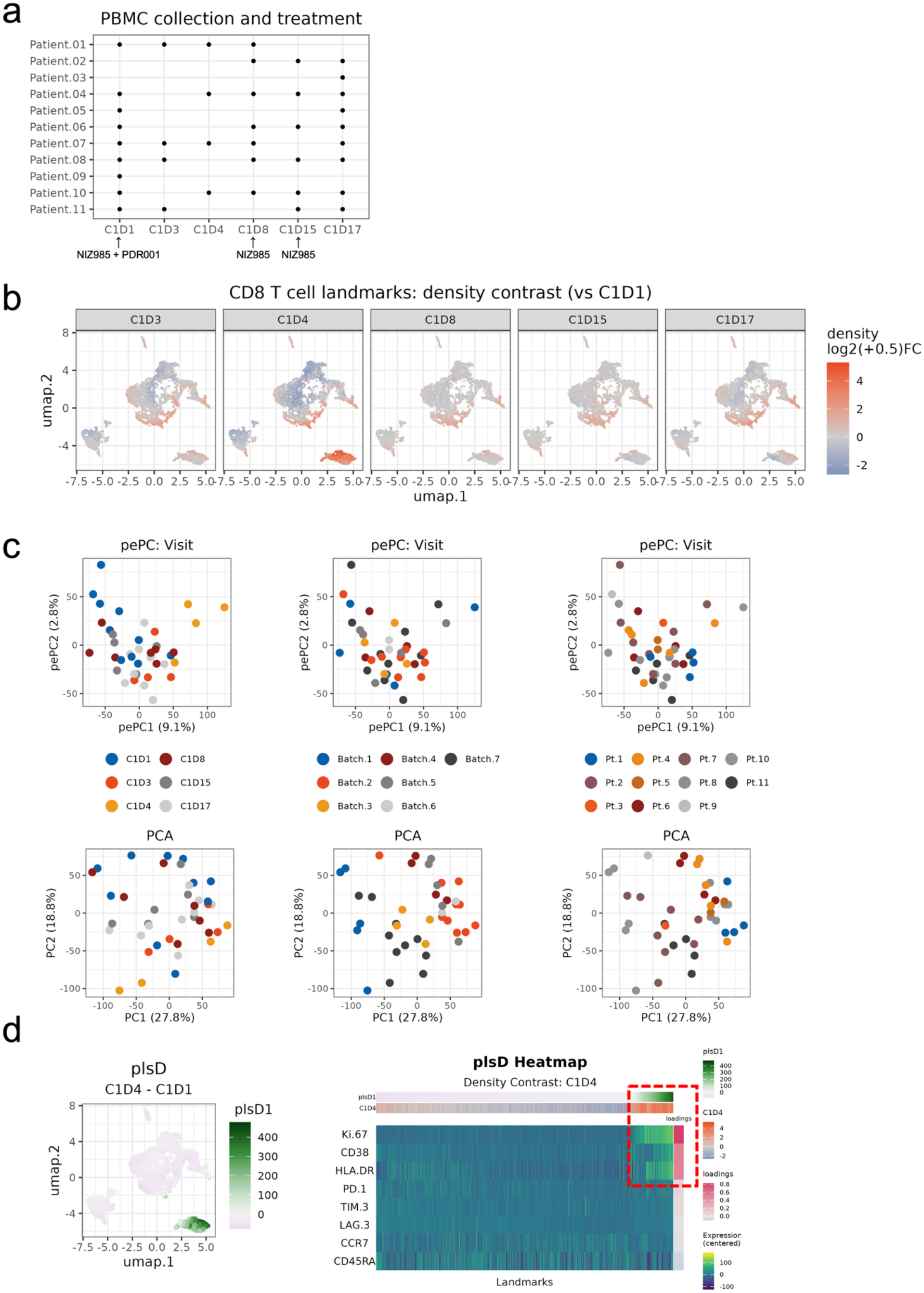
*tinydenseR* reveals treatment-responsive CD8 T cell states in an immuno-oncology clinical trial. (a) PBMC collection and treatment schedule across 11 patients. Dots indicate available samples for each patient at visits C1D1, C1D3, C1D4, C1D8, C1D15, and C1D17. Treatment annotations shown below the x-axis indicate NIZ985 + PDR001 at C1D1 and NIZ985 at C1D8 and C1D15. (b) UMAP projection of CD8 T cell landmarks colored by visit-specific density log2 fold-change relative to C1D1 for C1D3, C1D4, C1D8, C1D15, and C1D17. (c) Supervised (pePC; top row) and unsupervised (bottom row) sample embeddings colored by visit (left), batch (center), and patient ID (right). (d) Left: UMAP projection of CD8 T cell landmarks colored by plsD1 scores for the C1D4 vs C1D1 density contrast. Right: plsD heatmap for the same contrast, ordered by plsD1 score across landmarks. Annotation strips show plsD1 scores, the C1D4 vs C1D1 density contrast (both at the top), and plsD1 loadings (right-hand side of the heatmap). Rows show all markers used in the analysis, ordered by loadings.

**Figure S11.**
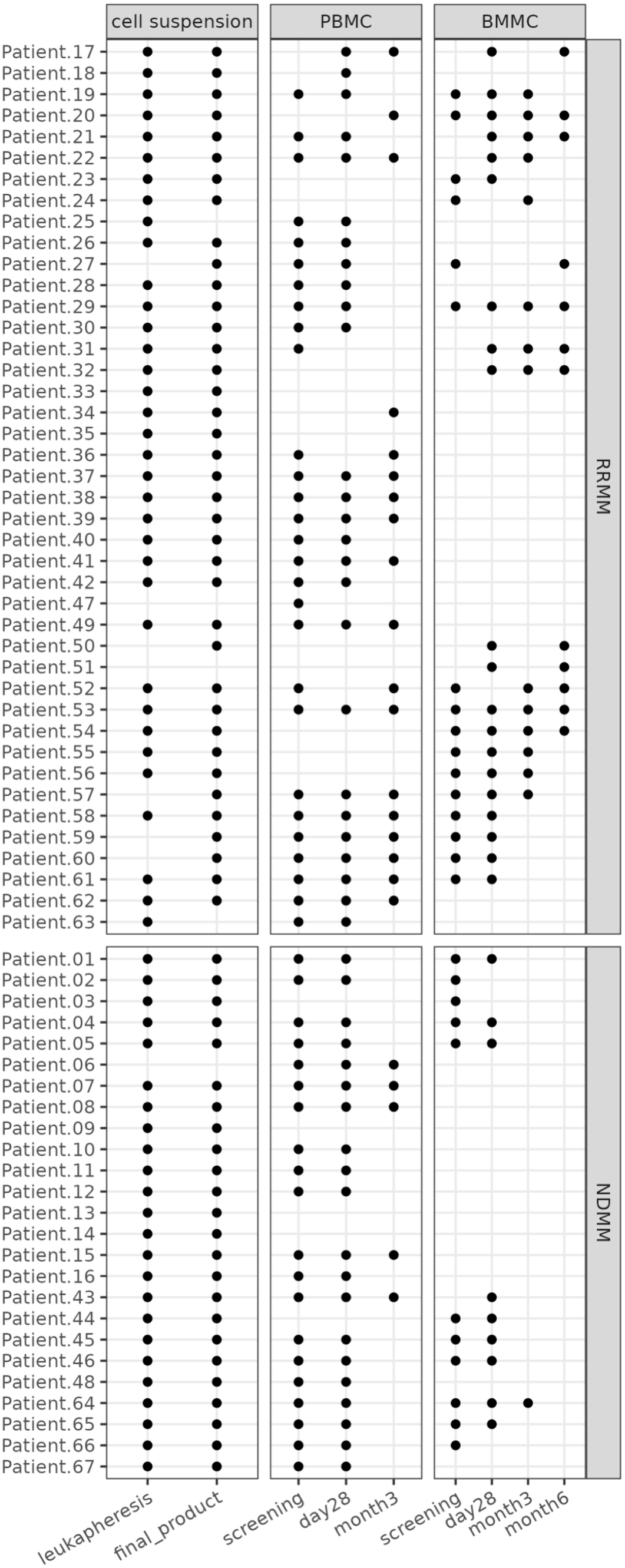
Sample availability across patients in the BCMA-targeting CAR T cell trial. Chart summarizing single-cell data availability across three compartments — cell suspension, peripheral blood mononuclear cells (PBMC), and bone marrow mononuclear cells (BMMC) — for patients enrolled in the phase 1 PHE885 CAR T trial. Patients are grouped by cohort (NDMM and RRMM), and time points include leukapheresis, final product, screening, day 28, month 3 and month 6 post-treatment. Black dots indicate availability of data for each patient and sample type.

**Figure S12.**
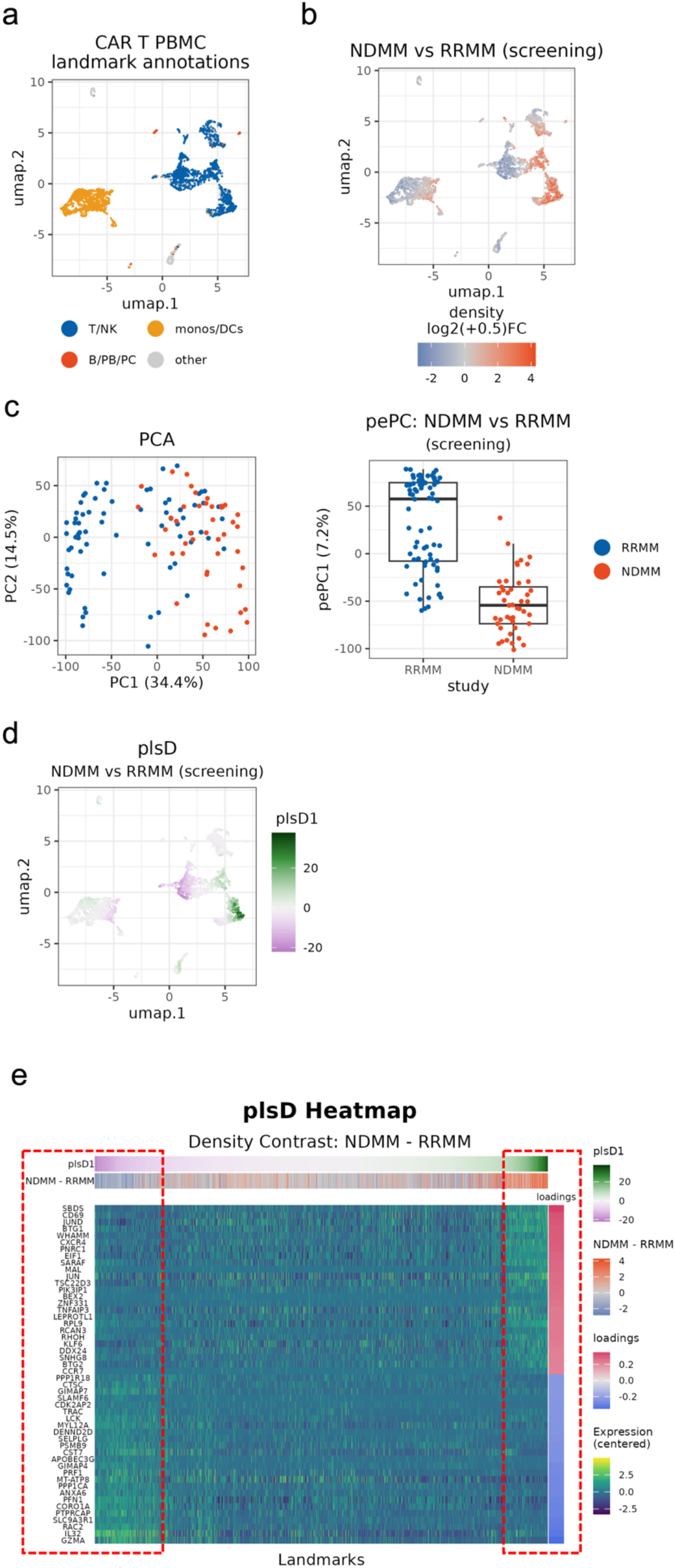
Modeling PBMC single-cell data at screening using *tinydenseR* reveals cohort-associated immune structure in NDMM and RRMM patients. (a) Landmark UMAP projection of peripheral blood mononuclear cells (PBMCs) colored by broad landmark annotations, showing T/NK, monocyte/dendritic-cell (monos/DCs), B/plasmablast/plasma-cell (B/PB/PC) and other landmark regions. (b) Landmark UMAP colored by the fitted density contrast for the screening NDMM versus RRMM comparison, displayed as density log2(+0.5) fold change. (c) Sample-level embedding computed from the log2-transformed landmark density matrix: unsupervised PCA (left) and supervised pePC for the screening NDMM-versus-RRMM contrast (right). (d) Landmark UMAP projection colored by plsD1 scores for the screening NDMM-versus-RRMM contrast. (e) plsD heatmap for the same contrast, showing the top 25 positively and negatively loaded features ranked by loading across the transcriptome.

**Figure S13.**
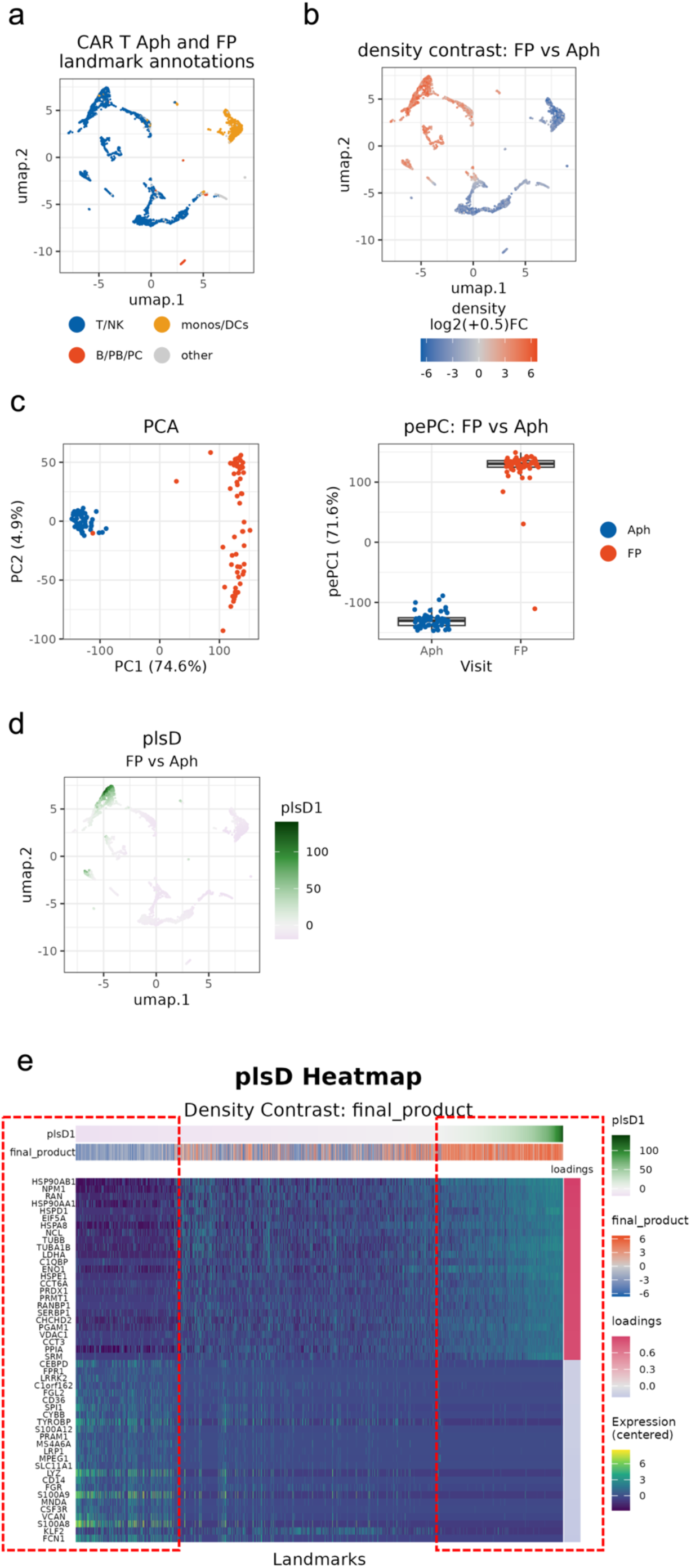
Modeling leukapheresis and final product single-cell data using *tinydenseR* reveals strong manufacturing-associated enrichment of activated T-cell states and depletion of non-T-cell contaminants. (a) Landmark UMAP projection of leukapheresis (Aph) and final product (FP) samples colored by broad landmark annotations, showing T/NK-, monocyte/dendritic-cell (monos/DCs)-, B/plasmablast/plasma-cell (B/PB/PC)- and other landmark regions. (b) Landmark UMAP colored by the fitted density contrast for FP versus Aph while controlling for cohort (NDMM or RRMM) and blocking for patient ID (right), displayed as density log2(+0.5) fold change. (c) Sample-level embedding computed from the log2-transformed landmark density matrix: unsupervised PCA (left) and supervised pePC1 for the FP versus Aph contrast. (d) Landmark UMAP projection colored by plsD1 scores for the FP versus Aph contrast. (e) plsD heatmap for the same contrast, showing the top 25 positively and negatively loaded features ranked by loading across the transcriptome.

## Notes

https://github.com/Novartis/tinydenseR

### Summary of Updates

We thank the reviewers for their careful and constructive evaluation. We agree that the original manuscript did not communicate the conceptual scope, novelty, or limitations of the framework as clearly as it should have. In response, we substantially revised both the presentation and the supporting analyses. The central revision reframes tinydenseR around a common landmark-by-sample fuzzy density matrix derived from UMAP-based cell-landmark connection strengths. This representation supports four downstream analysis modes: (1) landmark-level differential density modeling, (2) supervised quantitative sample embedding via partial-effect principal component projection (pePC), (3) density-contrast-aligned feature exploration via graph-diffused partial least squares decomposition (plsD), and (4) connection-strength-weighted pseudobulk differential expression. A dedicated algorithm overview ("The tinydenseR Algorithm") and revised Figure 1 make this structure explicit, and the revised manuscript clarifies which components are novel and which remain conventional once subsets are defined. To strengthen empirical support, we expanded synthetic and permutation benchmarks, added a landmark- versus cell-level integration comparison, updated miloR benchmarking to use graph refinement, and included analysis of the publicly available COMBAT COVID-19 PBMC dataset. The Methods now correct the terminology for UMAP-derived weights (connection strengths, not probabilities), make the landmark allocation rule explicit, and clarify assumptions underlying the density matrix construction. We do not claim that the revision resolves every benchmarking or sensitivity question raised. The goal was to make the claims more precise, the framework more transparent, and the empirical evidence more directly aligned with the revised conceptual framing.

## References

1. Cao, J., et al., The single-cell transcriptional landscape of mammalian organogenesis. Nature, 2019. 566(7745): p. 496–502.

2. Tabula Sapiens, C., et al., The Tabula Sapiens: A multiple-organ, single-cell transcriptomic atlas of humans. Science, 2022. 376(6594): p. eabl4896.

3. Regev, A., et al., The Human Cell Atlas. Elife, 2017. 6.

4. Han, X., et al., Construction of a human cell landscape at single-cell level. Nature, 2020. 581(7808): p. 303–309.

5. Liu, C., et al., Time-resolved systems immunology reveals a late juncture linked to fatal COVID-19. Cell, 2021. 184(7): p. 1836–1857 e22.

6. Sun, S., et al., Accuracy, robustness and scalability of dimensionality reduction methods for single-cell RNA-seq analysis. Genome Biol, 2019. 20(1): p. 269.

7. Wolf, F.A., P. Angerer, and F.J. Theis, SCANPY: large-scale single-cell gene expression data analysis. Genome Biol, 2018. 19(1): p. 15.

8. Butler, A., et al., Integrating single-cell transcriptomic data across different conditions, technologies, and species. 2018.

9. Luecken, M.D. and F.J. Theis, Current best practices in single-cell RNA-seq analysis: a tutorial. Mol Syst Biol, 2019. 15(6): p. e8746.

10. Kiselev, V.Y., T.S. Andrews, and M. Hemberg, Challenges in unsupervised clustering of single-cell RNA-seq data. Nat Rev Genet, 2019. 20(5): p. 273–282.

11. Lahnemann, D., et al., Eleven grand challenges in single-cell data science. Genome Biol, 2020. 21(1): p. 31.

12. Squair, J.W., et al., Confronting false discoveries in single-cell differential expression. Nat Commun, 2021. 12(1): p. 5692.

13. Tritschler, S., et al., Concepts and limitations for learning developmental trajectories from single cell genomics. Development, 2019. 146(12).

14. Nowicka, M., et al., CyTOF workflow: differential discovery in high-throughput high-dimensional cytometry datasets. F1000Res, 2017. 6: p. 748.

15. Weber, L.M., et al., diffcyt: Differential discovery in high-dimensional cytometry via high-resolution clustering. Commun Biol, 2019. 2: p. 183.

16. Amezquita, R., et al., Orchestrating Single-Cell Analysis with Bioconductor. 2020: Bioconductor. https://bioconductor.org/books/release/OSCA/

17. Hao, Y., et al., Dictionary learning for integrative, multimodal and scalable single-cell analysis. Nat Biotechnol, 2024. 42(2): p. 293–304.

18. Lun, A.T.L., A.C. Richard, and J.C. Marioni, Testing for differential abundance in mass cytometry data. Nat Methods, 2017. 14(7): p. 707–709.

19. Dann, E., et al., Differential abundance testing on single-cell data using k-nearest neighbor graphs. Nat Biotechnol, 2022. 40(2): p. 245–253.

20. Baran, Y., et al., MetaCell: analysis of single-cell RNA-seq data using K-nn graph partitions. Genome Biol, 2019. 20(1): p. 206.

21. Van Gassen, S., et al., FlowSOM: Using self-organizing maps for visualization and interpretation of cytometry data. Cytometry A, 2015. 87(7): p. 636–45.

22. Hie, B., et al., Geometric Sketching Compactly Summarizes the Single-Cell Transcriptomic Landscape. Cell Syst, 2019. 8(6): p. 483–493 e7.

23. Burkhardt, D.B., et al., Quantifying the effect of experimental perturbations at single-cell resolution. Nat Biotechnol, 2021. 39(5): p. 619–629.

24. Boyeau, P., et al., Deep generative modeling of sample-level heterogeneity in single-cell genomics. Nat Methods, 2025. 22(11): p. 2264–2274.

25. Ahlmann-Eltze, C. and W. Huber, Analysis of multi-condition single-cell data with latent embedding multivariate regression. Nat Genet, 2025. 57(3): p. 659–667.

26. COMBAT, A blood atlas of COVID-19 defines hallmarks of disease severity and specificity. Cell, 2022. 185(5): p. 916–938 e58.

27. Schulte-Schrepping, J., et al., Severe COVID-19 Is Marked by a Dysregulated Myeloid Cell Compartment. Cell, 2020. 182(6): p. 1419–1440 e23.

28. Amatangelo, M., et al., Pharmacodynamic changes in tumor and immune cells drive iberdomide’s clinical mechanisms of activity in relapsed and refractory multiple myeloma. Cell Rep Med, 2024. 5(6): p. 101571.

29. Dickinson, M.J., et al., A Novel Autologous CAR-T Therapy, YTB323, with Preserved T-cell Stemness Shows Enhanced CAR T-cell Efficacy in Preclinical and Early Clinical Development. Cancer Discov, 2023. 13(9): p. 1982–1997.

30. Jin, K., et al., An interactive single cell web portal identifies gene and cell networks in COVID-19 host responses. iScience, 2021. 24(10): p. 103115.

31. Yoshida, M., et al., Local and systemic responses to SARS-CoV-2 infection in children and adults. Nature, 2022. 602(7896): p. 321–327.

32. Parks, B. and W. Greenleaf, Scalable high-performance single cell data analysis with BPCells. bioRxiv, 2025.

33. Lun, A., BiocSingular: Singular Value Decomposition for Bioconductor Packages. 2025, Bioconductor.

34. Pagès, H., A. Lun, and P. Hickey, DelayedArray: A unified framework for working transparently with on-disk and in-memory array-like datasets. 2025, Bioconductor.

35. Alice, K., J.C. Marioni, and M.D. Morgan, 2023.

36. Kang, J.B., et al., Efficient and precise single-cell reference atlas mapping with Symphony. Nat Commun, 2021. 12(1): p. 5890.

37. Stuart, T., et al., Comprehensive Integration of Single-Cell Data. Cell, 2019. 177(7): p. 1888–1902 e21.

38. Hao, Y., et al., Integrated analysis of multimodal single-cell data. Cell, 2021. 184(13): p. 3573–3587 e29.

39. Joodaki, M., et al., Detection of PatIent-Level distances from single cell genomics and pathomics data with Optimal Transport (PILOT). Mol Syst Biol, 2024. 20(2): p. 57–74.

40. De Donno, C., et al., Population-level integration of single-cell datasets enables multi-scale analysis across samples. Nat Methods, 2023. 20(11): p. 1683–1692.

41. Willem, T., et al., Biases in machine-learning models of human single-cell data. Nat Cell Biol, 2025. 27(3): p. 384–392.

42. Persad, S., et al., SEACells infers transcriptional and epigenomic cellular states from single-cell genomics data. Nat Biotechnol, 2023. 41(12): p. 1746–1757.

43. Skinnider, M.A., et al., Cell type prioritization in single-cell data. Nat Biotechnol, 2021. 39(1): p. 30–34.

44. Nadig, A., et al., Transcriptome-wide analysis of differential expression in perturbation atlases. Nat Genet, 2025. 57(5): p. 1228–1237.

45. Hafner, L., et al., Single-cell differential expression analysis between conditions within nested settings. Brief Bioinform, 2025. 26(4).

46. Drineas, P., et al., Fast Approximation of Matrix Coherence and Statistical Leverage. Journal of Machine Learning Research, 2012. 13: p. 3475–3506.

47. Baglama, J. and L. Reichel, Augmented implicitly restarted Lanczos bidiagonalization methods. Siam Journal on Scientific Computing, 2005. 27(1): p. 19–42.

48. Korsunsky, I., et al., Fast, sensitive and accurate integration of single-cell data with Harmony. Nat Methods, 2019. 16(12): p. 1289–1296.

49. McInnes, L., J. Healy, and J. Melville, UMAP: Uniform Manifold Approximation and Projection for Dimension Reduction. arXiv, 2020: p. 1802.03426.

50. Csárdi, G. and T. Nepusz, The igraph software package for complex network research. InterJournal, 2006. Complex Systems: p. 1695.

51. Traag, V.A., L. Waltman, and N.J. van Eck, From Louvain to Leiden: guaranteeing well-connected communities. Sci Rep, 2019. 9(1): p. 5233.

52. Stoeckius, M., et al., Simultaneous epitope and transcriptome measurement in single cells. Nat Methods, 2017. 14(9): p. 865–868.

53. Ritchie, M.E., et al., limma powers differential expression analyses for RNA-sequencing and microarray studies. Nucleic Acids Res, 2015. 43(7): p. e47.

54. Phipson, B., et al., Robust Hyperparameter Estimation Protects against Hypervariable Genes and Improves Power to Detect Differential Expression. Ann Appl Stat, 2016. 10(2): p. 946–963.

55. Ignatiadis, N., et al., Data-driven hypothesis weighting increases detection power in genome-scale multiple testing. Nat Methods, 2016. 13(7): p. 577–80.

56. Boca, S.M. and J.T. Leek, A direct approach to estimating false discovery rates conditional on covariates. PeerJ, 2018. 6: p. e6035.

57. Chen, Y., et al., edgeR v4: powerful differential analysis of sequencing data with expanded functionality and improved support for small counts and larger datasets. Nucleic Acids Res, 2025. 53(2).

58. Hanzelmann, S., R. Castelo, and J. Guinney, GSVA: gene set variation analysis for microarray and RNA-seq data. BMC Bioinformatics, 2013. 14: p. 7.

59. Dolgalev, I., msigdbr: MSigDB Gene Sets for Multiple Organisms in a Tidy Data Format. 2025, CRAN.

60. Angerer, P., et al., destiny: diffusion maps for large-scale single-cell data in R. Bioinformatics, 2016. 32(8): p. 1241–3.

61. Wickham, H., et al., profvis: Interactive Visualizations for Profiling R Code. 2024, CRAN.

62. Finak, G. and M. Jiang, flowWorkspace: Infrastructure for representing and interacting with gated and ungated cytometry data sets. 2025, Bioconductor.

63. Young, M.D. and S. Behjati, SoupX removes ambient RNA contamination from droplet-based single-cell RNA sequencing data. Gigascience, 2020. 9(12).

64. Hafemeister, C. and R. Satija, Normalization and variance stabilization of single-cell RNA-seq data using regularized negative binomial regression. Genome Biol, 2019. 20(1): p. 296.

## Supplemental References

65. R Core Team, R: A Language and Environment for Statistical Computing. 2025, R Foundation for Statistical Computing.

66. Huber, W., et al., Orchestrating high-throughput genomic analysis with Bioconductor. Nature Methods, 2015. 12(2): p. 115–21.

67. Krantz, S., collapse: Advanced and Fast Statistical Computing and Data Transformation in R. 2024, arXiv.

68. Wickham, H., et al., dplyr: A Grammar of Data Manipulation. 2023, CRAN.

69. Ooms, J., curl: A Modern and Flexible Web Client for R. 2025.

70. Saelens, W., et al., A comparison of single-cell trajectory inference methods. Nat Biotechnol, 2019. 37(5): p. 547–554.

71. Wickham, H., M. Kuhn, and D. Vaughan, generics: Common S3 Generics not Provided by Base R Methods Related to Model Fitting. 2025, CRAN.

72. Korotkevich, G., et al., Fast gene set enrichment analysis. bioRxiv, 2021.

73. Arora, S., et al., GenomeInfoDb: Utilities for manipulating chromosome names, including modifying them to follow a particular naming style. 2025, Bioconductor.

74. Lawrence, M., et al., Software for computing and annotating genomic ranges. PLoS Comput Biol, 2013. 9(8): p. e1003118.

75. van den Brand, T., ggh4x: Hacks for ‘ggplot2’. 2025.

76. Campitelli, E., ggnewscale: Multiple Fill and Colour Scales in ‘ggplot2’. 2025.

77. Wickham, H., ggplot2: Elegant Graphics for Data Analysis. 2016, CRAN.

78. Slowikowski, K., ggrepel: Automatically Position Non-Overlapping Text Labels with ‘ggplot2’. 2026, CRAN.

79. Auguie, B., gridExtra: Miscellaneous Functions for “Grid” Graphics. 2017.

80. Wickham, H. and T.L. Pedersen, gtable: Arrange ‘Grobs’ in Tables. 2024, CRAN.

81. Korsunsky, I., et al., harmony: Fast, Sensitive, and Accurate Integration of Single Cell Data. 2024.

82. Csárdi, G., et al., igraph: Network Analysis and Visualization in R. 2025, CRAN.

83. Baglama, J., L. Reichel, and B.W. Lewis, irlba: Fast Truncated Singular Value Decomposition and Principal Components Analysis for Large Dense and Sparse Matrices. 2022.

84. Xie, Y., knitr: A General-Purpose Package for Dynamic Report Generation in R. 2025, CRAN.

85. Bates, D., M. Maechler, and M. Jagan, Matrix: Sparse and Dense Matrix Classes and Methods. 2025, CRAN.

86. Ahlmann-Eltze, C., P. Hickey, and H. Pagès, MatrixGenerics: S4 Generic Summary Statistic Functions that Operate on Matrix-Like Objects. 2025, Bioconductor.

87. Bengtsson, H., matrixStats: Functions that Apply to Rows and Columns of Matrices (and to Vectors). 2025.

88. Pedersen Thomas, L., patchwork: The Composer of Plots. 2025.

89. Kolde, R., pheatmap: Pretty Heatmaps. 2025.

90. Storey John, D., et al., qvalue: Q-value estimation for false discovery rate control. 2025.

91. Eddelbuettel, D., Seamless R and C++ Integration with Rcpp. 2013

92. Junji, N. and E.-j. Nakama, RhpcBLASctl: Control the Number of Threads on ‘BLAS’. 2023.

93. Henry, L. and H. Wickham, rlang: Functions for Base Types and Core R and ‘Tidyverse’ Features. 2025, CRAN.

94. Pagès, H., M. Lawrence, and P. Aboyoun, S4Vectors: Foundation of vector-like and list-like containers in Bioconductor. 2025, Bioconductor.

95. Wickham, H., L. Pedersen Thomas, and D. Seidel, scales: Scale Functions for Visualization. 2025, CRAN.

96. Kulichova, T. and M. Kratochvil, scattermore: Scatterplots with More Points. 2023.

97. Pagès, H., Seqinfo: A simple S4 class for storing basic information about a collection of genomic sequences. 2025, bioconductor.

98. Hoffman, P., et al., SeuratObject: Data Structures for Single Cell Data. 2025, CRAN.

99. Pebesma Edzer, J. and R. Bivand, Classes and methods for spatial data in R. 2005, CRAN.

100. Ahlmann-Eltze, C., sparseMatrixStats: Summary Statistics for Rows and Columns of Sparse Matrices. 2025, Bioconductor.

101. Morgan, M., et al., SummarizedExperiment: A container (S4 class) for matrix-like assays. 2025, Bioconductor.

102. Leek Jeffrey, T., et al., swfdr: Estimation of the science-wise false discovery rate and the false discovery rate conditional on covariates. 2025.

103. Kang, J., I. Korsunsky, and S. Raychaudhuri, symphony: Efficient and Precise Single-Cell Reference Atlas Mapping. 2023.

104. Wickham, H., D. Vaughan, and M. Girlich, tidyr: Tidy Messy Data. 2024, CRAN.

105. Henry, L. and H. Wickham, tidyselect: Select from a Set of Strings. 2024, CRAN.

106. Melville, J., uwot: The Uniform Manifold Approximation and Projection (UMAP) Method for Dimensionality Reduction. 2025, CRAN.

